# Multi-dimensional analyses identify genes of high priority for pancreatic cancer research

**DOI:** 10.1101/2021.05.28.446056

**Authors:** Zeribe C. Nwosu, Heather Giza, Maya Nassif, Verodia Charlestin, Rosa E. Menjivar, Daeho Kim, Samantha B. Kemp, Nina G. Steele, Jiantao Hu, Biao Hu, Shaomeng Wang, Marina Pasca di Magliano, Costas A. Lyssiotis

## Abstract

Genomic profiling has unveiled the molecular subtypes and mutational landscape of pancreatic ductal adenocarcinoma (PDAC). However, there is a knowledge gap on the consistency of gene expression across PDAC tumors profiled in independent studies and this limits follow up research. To facilitate novel drug target prioritization and biomarker discovery, we investigated the most consistently expressed genes in human PDAC. We identified ~4,000 genes highly or lowly expressed in at least 4 of 5 microarrays (adjusted *P*<0.05) and validated their expression pattern in additional datasets, bulk tumor and single-cell RNA sequencing samples. Over 50% of the genes were previously uncharacterized in PDAC; many correlated with proliferation, metastasis, mutation, tumor grade, and ~41% predicted overall survival. We identified 185 high-priority targets (notably in cell cycle and glycolysis) whose inhibition suppressed PDAC cell viability in multiple RNA interference datasets and these genes predicted treatment in mouse models. Our results represent important milestone in the quest for mechanisms, drug targets and biomarkers in PDAC, and originate from an adaptable analytical concept that can aid discovery in other cancers.

**Highlights:**

- Identifies ~4,000 consistent genes across PDAC microarrays, >50% of which have not been studied
- Glycolysis and cell cycle are the most consistent processes in PDAC
- Heterogeneous pathways underlie or correlate with clinicopathological variables
- Identifies 205 genes with similar expression pattern in PDAC tissues and peripheral blood
- Highlights 185 upregulated genes that are high priority therapeutic targets in PDAC

---

Pancreatic ductal adenocarcinoma (PDAC) is an aggressive, highly metastatic and drug resistant disease, and as a result is one of the most lethal solid tumors^1^. Prior studies have unanimously converged on *KRAS, TP53, SMAD4* and *CDKN2A* as the most frequently mutated genes in PDAC^2–5^. These oncogenic mutations, notably KRAS, dictate drug sensitivity, drive tumor initiation and facilitate progression^6,7^. Studies of gene expression profiles in PDAC have defined two to four major subtypes of PDAC^2–4,8–10^; these subtypes also impact therapeutic response and patient overall survival. Several other studies have performed gene expression profiling, RNA sequencing, proteomics, or metabolomics towards unravelling the unique molecular features that distinguish PDAC tissues from healthy pancreatic tissues. These multi-omics approaches facilitate hypothesis generation and novel discoveries on the mechanisms and therapeutic opportunities in PDAC.

Although studies have generated an extensive amount of gene expression data from PDAC, genes that are frequently lowly or highly expressed in the tumors from the multiple datasets are largely unknown. Experimental validation studies often rely on data from small patient cohorts, most times on as few as one dataset, and focus on specific genes/pathways. This approach underestimates the scope of PDAC alterations, and several of those validated genes are not reproducibly changed in independent datasets. Other issues such as the actual proportion of PDAC cells in a tumor mass as well as sampling (e.g., surgically resectable vs non-resectable tumors) limit the clinical utility of gene findings. Establishing the consistent genes in PDAC and their expression pattern will facilitate drug target identification and prioritization, aid biomarker discovery, and enhance the prospects of finding the core molecular mechanisms driving this intractable disease.

In this study, we first analyzed five published PDAC microarrays and defined a gene as ‘consistent’ if it is ‘upregulated’ or ‘downregulated’ in at least 4 cohorts (adjusted *P*<0.05). We followed up with analyses and cross-validation of these genes in over 20 datasets, including microarrays (tumors and blood); bulk tumor and single cell RNA sequencing; normal tissues and cancer cell line gene expression; drug response, and RNA interference datasets. We interrogated the association of the genes with several variables such as tumor subtypes, proliferation, metastasis, mutation and tumor grade, using at least two patient datasets in each case. Further, our analysis of Kaplan-Meier overall survival, Cox proportional hazards regression and gene essentiality (four datasets in each case) is to date the most extensive assessment of gene consistency relative to their potential clinical relevance. In cell lines and mouse models, we demonstrate the utility of the consistent genes in predicting drug response. Together, our findings will facilitate gene prioritization for basic, translational and clinical PDAC research, and thus pave way for better treatment opportunities for this disease.

## Results

### Consistently expressed genes in PDAC tissues

To identify the most consistently expressed and potentially important genes in PDAC, we analyzed over 20 publicly accessible datasets of gene expression or RNA interference (**Fig. 1a, Extended Data Fig. 1a**). We started with five microarray datasets containing a total of 331 pancreatic tumors and 204 non-tumoral tissue samples (**Extended Data Fig. 1b**). We described a gene as ‘consistent’ if its expression is high (or low) in at least 4 of the 5 datasets. Based on this criterion, we identified 2,010 consistently upregulated genes (hereafter CUGs) and 1,928 consistently downregulated genes (CDGs) (adjusted *P*<0.05). As many as 993 (49% of CUGs) and 737 (38% of CDGs) were consistent in all five datasets (**Fig. 1a**), indicating that these genes have reproducible expression pattern in PDAC. The topmost 20 CUGs included *LAMC2, TMPRSS4, S100P, SLC6A14, COL10A1, CTSE, LAMB3, CEACAM5/6*, and glucose transporter *SLC2A1* (**Fig. 1b**), and these appeared within the top 1-5% of highly expressed genes in all five datasets (**Supplementary Table 1a**). On the other hand, the lowest CDGs included *AOX1, IAPP*, albumin (*ALB*), *SERPINI2, PDK4* and *PNLIPRP1/2* (**Fig. 1b**) and were also consistently the lowest expressed across the patient datasets.

**Figure. 1.**
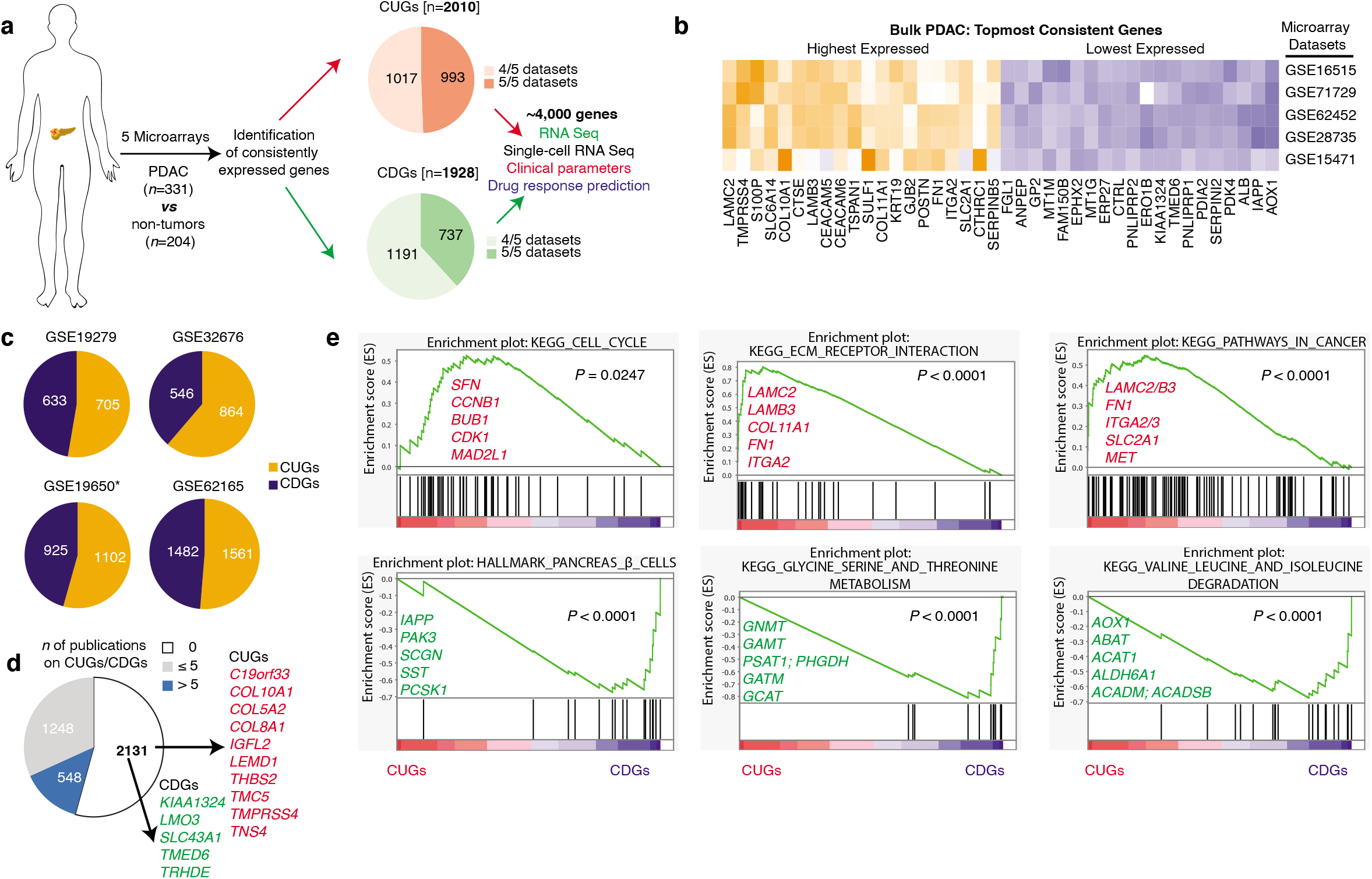
Consistently expressed genes in human PDAC tissues. **a,** Schematic overview illustrating the identification and potential utility of the consistent genes. Five PDAC microarray datasets were used for the identification of the genes (see Fig. 1b) and ‘consistent’ was defined as genes significantly up or downregulated in at least 4 datasets (adjusted *P*<0.05). CUGs – consistently upregulated genes; CDGs – consistently downregulated genes. See Methods and Extended Data Fig. 1b for sample sizes. **b**, Topmost 20 highly and lowly expressed genes (i.e., CUGs and CDGs respectively) based on average expression rank in the five datasets. **c**, Number of CUGs and CDGs with the same high or low pattern in the independent microarray datasets GSE19279, GSE32676, GSE19650 and GSE62165. *Dataset of pre-malignant tumor stages. See Methods for sample sizes/types. **d**, Pie chart indicating the novelty of the consistent genes (i.e., CUGs and CDGs) in PDAC based on the number (*n*) of prior publications (0, ≤5 and >5) as observed via PubMed search. Highlighted in red are topmost CUGs with no prior publication; in green are topmost CDGs with no prior publication. **e**, Gene set enrichment plots of KEGG pathways or Hallmark associated with the consistent genes. The plots were generated using the average ranked expression score of the consistent genes (*n*=3,938) that are significantly changed at adjusted *P*<0.05 at least in 4 of the 5 ‘discovery’ datasets.

We analyzed the expression of the CUGs and CDGs in four independent PDAC patient cohorts and found that between 28% to ~80% of CUGs and CDGs were also high and low, respectively (**Fig. 1c, Supplementary Note 1**). Further, the topmost genes in the initial five datasets also appeared topmost in those four independent datasets (**Extended Data Fig. 1c**) supporting that CUGs and CDGs are expected to be reproducibly changed in most PDAC cohorts.

In the initial five datasets, we also identified 2,722 additional genes (*n*=1,063 up- and 1,659 down) with consistent expression pattern in three of the five datasets (adjusted *P*<0.05) (**Supplementary Table 1b**). These included well-known genes in cancer such as *ALDOC, HIF1A* (in glycolysis), *SNAI2, ZEB1* (in epithelial-mesenchymal transition), monocarboxylate transport 4 (*SLC16A3*), *MMP2, GPNMB, KRT17* and *INHBA* (all upregulated) as well as downregulated genes such as *GATA4, ADH1A, PTF1A, CBS*, and genes encoding pancreatic digestive enzymes (e.g., *PRSS1, CPA1* and *CPB1*) (**Extended Data Fig. 1d, Supplementary Table 1b**). Several of these genes, though not emphasized in our analysis, also maintained similar expression pattern in the four independent cohorts, altogether revealing an extremely complex molecular portrait of PDAC. As expected, many CUGs and CDGs have been mechanistically studied or proposed as PDAC biomarkers or therapeutic targets. However, we found no substantial prior PDAC publication on >50% of the ~4000 consistent genes (**Fig. 1d**, **Supplementary Table 2**), indicating the enormity of untapped opportunities for new discoveries.

To determine if the consistent genes reflect typical cancer profiles, we performed pathway enrichment and gene ontology (GO) analyses. The ‘cell cycle’, ‘extracellular matrix receptor interaction’, ‘pathways in cancer’, ‘focal adhesion’, ‘p53’, and ‘transcription factor binding’ emerged among the upregulated processes and are consistent with oncogenic alterations. These were accompanied by evidence of disrupted pancreatic beta cell homeostasis and profound metabolic alterations (**Fig. 1e, Extended Data Fig. 1e**). Interactome and disease ontology analyses further associated the consistent genes with precancerous conditions, PDAC and other malignancies (**Extended Data Fig. 1f-g**). Altogether, we have uncovered the most consistently upregulated and suppressed genes in human PDAC tissues and note an overwhelming paucity of experimental data on the role of these genes in this cancer type.

### Multiple molecular pathways underlie PDAC

We examined specific pathways for the presence of the consistently deregulated genes. We focused on metabolism/transport, signaling, immunity, cell cycle and transcriptional regulation/epigenetics, all of which appeared in the pathway analysis – with an additional focus on transporters of which most are novel in PDAC. The number of genes we identified per process ranged from 45 in cell cycle, 76 in immunity, 229 transporters, 459 in metabolism, 549 in signaling to 723 transcription factors and 32 epigenetic components (**Supplementary Table 3**). Across the PDAC cohorts, most cell cycle genes (39 out of 45) were upregulated (i.e., CUGs), topmost of which are *SFN, CCNB1, BUB1, CDK1*, and *MAD2L1*. The remaining six were CDGs namely *ANAPC13, ANAPC5, MAD2L2, GADD45G, CCND2* and surprisingly *MYC* (**Fig. 2a**), which has been extensively implicated as a driver of cancer. In immunity, 60 of the 76 genes were CUGs, most notable of which were *CXCL5, ITGA3, CCL20, IFI44L*, and *IFI27* (an unknown gene in PDAC), and the most enriched immune hallmarks were interferon alpha and gamma, TNFA/NF-κB, ‘inflammatory response’ and ‘IL2/STAT5’signaling (**Fig. 2b, Extended Data Fig. 2a**).

**Figure 2.**
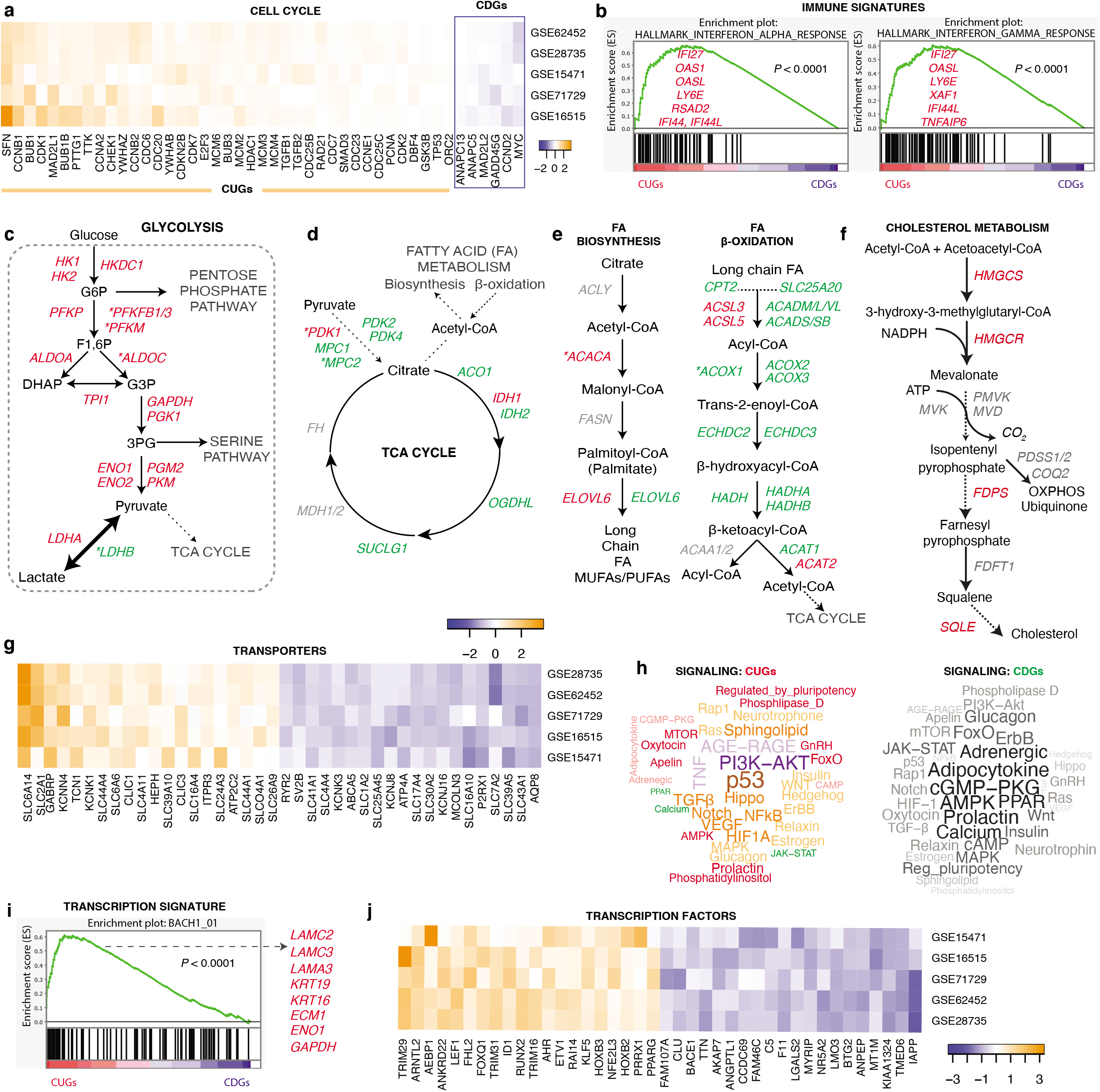
The consistent genes are components of various pathways. **a,** Components of cell cycle that showed consistent expression pattern. **b,** Gene set enrichment plots indicating upregulated consistent immune signatures. The indicated genes are top in the pathways shown. **c,** Schematic depiction of the genes in the glycolytic pathway. **d**, Depiction of genes in the tricarboxylic acid (TCA) cycle. **e**, Depiction of genes in fatty acid biosynthesis and breakdown (beta oxidation). **f**, Depiction of genes in cholesterol metabolism. **g,** Heatmap showing the most consistently expressed nutrient and ion transporters (top 20 up/downregulated). **h,** Word cloud showing signaling processes associated with the CUGs and CDGs. **i,** Gene set enrichment plot of transcriptional signatures, indicating upregulated BACH1_01 signatures. Highlighted genes (in red) were among the topmost of enriched signatures of BACH1_01 signatures. **j,** Heatmap showing the most consistently expressed transcription factors (top 20 up/downregulated). **Fig. 2c-f**, * indicates genes identified as consistent in 3/5 datasets; all other indicated genes have consistent expression pattern in ≥4 datasets; in red – CUGs, in green – CDGs, in gray – genes not captured as consistent and are shown given their known position/role in the respective pathways.

In metabolism, glycolysis genes emerged as the most consistently upregulated in PDAC (**Fig. 2c**). Other notably upregulated metabolic genes included *SULF1/2* (in glycan metabolism), *GPX2/8* (in glutathione metabolism), *NQO1* and *NOX4* (in oxidation/reduction reaction) (**Supplementary Table 3**). In contrast, the serine biosynthetic pathway, which branches off from glycolysis, was down (**Extended Data Fig. 2b).** Most components of the tricarboxylic acid (TCA) cycle, which connects glycolysis to oxidative phosphorylation (OXPHOS), were down or not consistent (**Fig. 2d**). The OXPHOS ‘hallmark’ also emerged low (**Extended Data Fig. 2c**) as were components of fatty acid beta oxidation (**Fig. 2e**), which altogether reflect a potential vulnerability in PDAC mitochondrial metabolism. PDAC showed a selective upregulation of genes in fatty acid biosynthesis and cholesterol metabolism (**Fig. 2e-f**). Many genes in transsulfuration pathway and amino acid metabolism were low, with a notable exception of tryptophan metabolism, where *TDO2* and *KYNU* (both CUGs) were identified (**Extended Data Fig. 2c**). Thus, although metabolism is generally considered altered in PDAC, these analyses reveal specific pathway components that their expression are more likely correlated with carcinogenesis.

Of the 229 transporters, the neurotransmitter/amino acid transporter (*SLC6A14*), glucose transporter (*SLC2A1*), *GABRP, KCNN4*, and cobalamin transporter (*TCN1*) were topmost CUGs, whereas *P2RX1, SLC7A2, SLC39A5, SLC43A1* and aquaporin 8 (*AQP8*) are the lowest CDGs (**Fig. 2g)**. With respect to signaling, we analyzed genes across 39 major signaling pathways in the KEGG database. Known processes such as HIF-1, p53, MAPK, Hippo, and AGE-RAGE signaling emerged as highly upregulated (**Fig. 2h**). In contrast, components of cGMP-PKG, PPAR and AMPK signaling were mostly downregulated (**Fig. 2h**). Of note, virtually every pathway had genes consistently up- and downregulated and implication of such dichotomous expression pattern is unknown. Regarding transcriptional signatures, BACH1 was the top enriched (**Fig. 2i)** and was recently reported to drive metastasis and the expression of glycolysis genes (*HK2* and *GAPDH*) in lung cancer^11,12^. On the other hand, AR and HNF1 transcriptional signatures are suppressed in PDAC (**Extended Data Fig. 2d**). We found that topmost upregulated transcription factors are *TRIM29, ARNTL2, AEBP1 ANKRD22* and *LEF1* (in Wnt signaling), whereas the lowest expressed are *IAPP, TMED6, KIAA1324. MT1M* and *ANPEP* (**Fig. 2j**). Several epigenetic targets also emerged, including *KDM5B, EZH2, HDAC1, HELLS, DNMT1, KDM2A/1B* and histone protein encoding genes *HIST1H2BC, HIST1H2BD, HIST1H2BE, HIST1H4H, HIST2H2BE* (all CUGs), whereas *SIRT3, CXXC1, ING3, KDM8, CITED2/4, KAT2B* topped the downregulated epigenetic targets in PDAC (**Extended Data Fig. 2e)**. These data shed light on the extensive molecular complexity of PDAC and highlight specific components that could be prime targets for mechanistic studies.

### The consistent genes show tumor-microenvironment cell expression pattern and are differentially expressed in blood samples

Tumor-stromal interactions are important in PDAC initiation and maintenance and have emerged as a focus area for PDAC therapy^10,13–15^. Previous studies aimed to distinguish neoplastic from stromal signatures via microdissection^10,14^. We applied a different approach by first defining the consistent genes in tumors compared to non-tumors, and then using these genes to further determine the expression of the consistent genes in normal pancreas relative to other normal tissues using three datasets (**Fig. 3a, Extended Data Fig. 3a-c**). We expected a tumor-specific CUGs to be low in normal pancreas and, reciprocally, tumor-specific CDGs to be high – a criterion that should accurately refine consistent genes that are potential therapeutic targets or altered at the early tumor stage. Indeed, 66% (*n*=1,333) of the CUGs and 72% (*n*=1,384) of CDGs met these tumor-specific criteria in ≥2 datasets (**Fig. 3b-3c**). Clustering analyses showed that glycolytic genes *HK1*, *GAPDH, ENO1*, *LDHA*, and *PKM*, as well as cell cycle genes (except *CCND2*) were among the lowest expressed CUGs in normal pancreas. Others included *TSC22D1, B2M, S100A6* and *CFL1*. For the CDGs, *CEL, GP2, REG1A, REG1B, REG3G, PRSS3, PNLIPRP1* and *PLA2G1B* were among the highest expressed in normal pancreas (**Extended Data Fig. 3a-c, Supplementary Table 4a).** We noticed these CDGs were highly expressed in the ‘exocrine-like’ tumor subtype defined by Collison et al^3^, suggesting that tissues in that subtype were mainly normal or well-differentiated.

**Figure 3.**
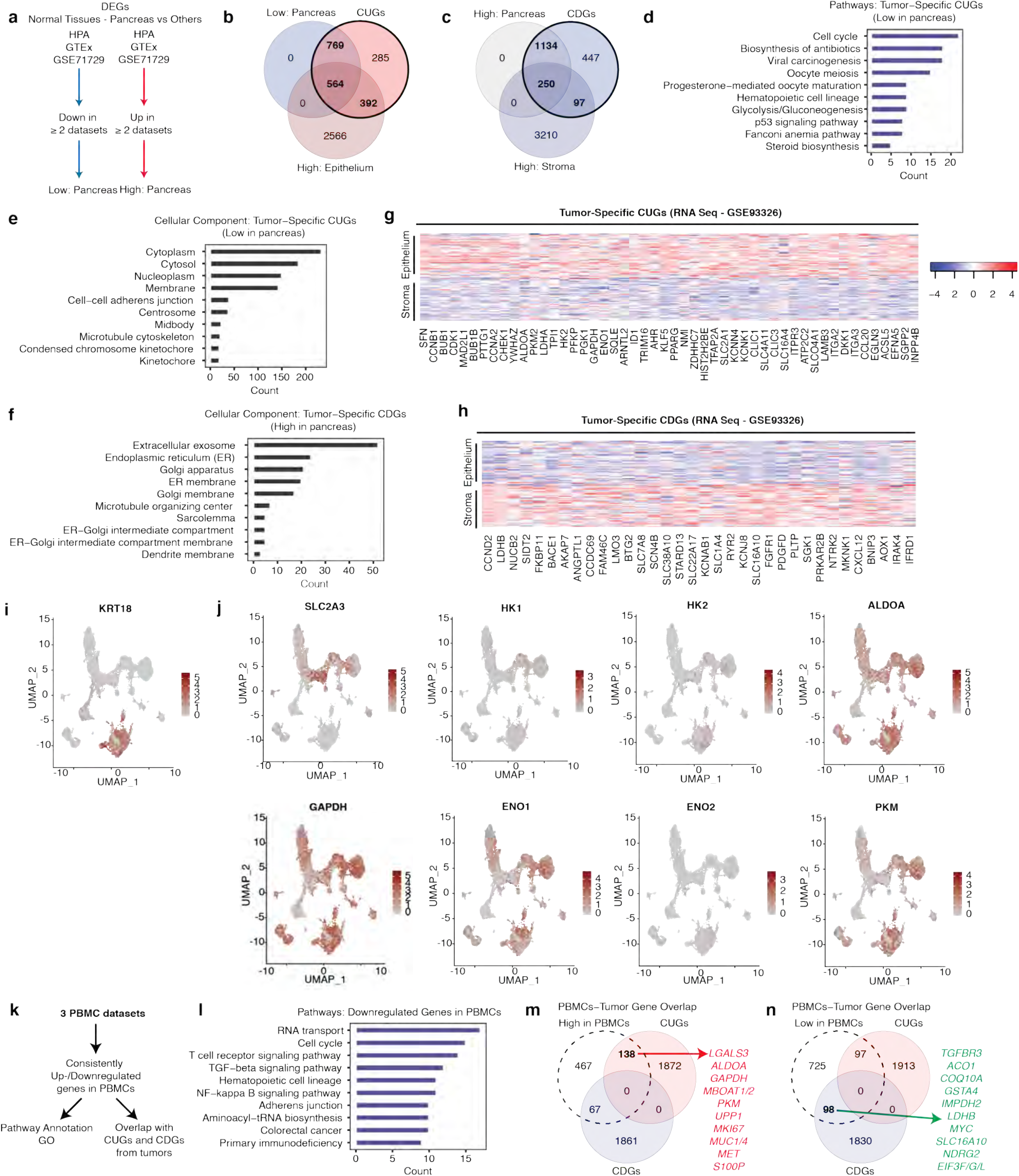
Consistent genes show tumor-specificity and expression changes in patients’ blood. **a,** Workflow for the identification of tumor-microenvironmental cell expressed CUGs and CDGs. Expression in PDAC was compared to normal tissues in the Human Protein Atlas (HPA, *n*=36 tissue types), Genotype-Tissue Expression (GTEx, *n=36* tissue types) RNA seq data and GSE71729 dataset (normal tissues: pancreas, *n*=46; liver, *n*=27; lung, *n*=19; lymph node, *n*=10; spleen, *n*=11). Similar expression pattern in ≥2 of HPA, GTEx or GSE71729 was the gene selection criteria. DEGs – differentially expressed genes. **b,** Venn diagram showing CUGs overlap in normal pancreas and with highly expressed genes in epithelium compared to stroma from laser microdissection dataset (GSE93326). **c,** Venn diagram showing CDGs overlap in normal pancreas and with highly expressed genes in the stroma (low in epithelium). **d,** Pathway annotation of the 564 tumor-specific CUGs (high in the epithelium and lowly expressed in normal pancreas). **e,** GO cellular component associated with the 564 tumor-specific CUGs (high in the epithelium and lowly expressed in normal pancreas). **f,** GO cellular component associated with the 250 tumor-specific CDGs (low in the epithelium relative to stroma and highly expressed in normal pancreas). Bars in Figure 3e-f are colored ‘darkgray’ to distinguish the plots (from normal pancreas) for that of cancer tissues. **g,** Heatmap showing differential expression of the topmost tumor-specific CUGs in the tumor epithelium. **h,** Heatmap showing differential expression of the lowest tumor-specific CDGs in the tumor epithelium. **i**, Uniform manifold approximation and projection (UMAP) of single cell RNA sequencing data depicting epithelial and immune cell population as marked by *KRT19* and *PTPRC* expression, respectively. **j**, UMAP showing the expression of glycolysis genes in the epithelial, immune cells or in both compartments. **k**, Schematic overview of the analysis of peripheral blood mononuclear cell (PBMCs) datasets (GSE15932, GSE49641, and GSE74629, *P*<0.05). **l**, Pathway analysis of consistently downregulated genes in 2 of GSE15932, GSE49641, and GSE74629 PBMC datasets. **m-n**, Venn diagram showing overlapping genes between CUGs and CDGs from tissues and consistent genes in PBMCs.

To further verify tumor-specificity, we overlapped the normal pancreas genes from the human protein atlas, genotype-tissue expression and or GSE71729 datasets with the laser capture microdissection (LCM) RNA seq data of PDAC epithelium versus stromal cells (GSE93326, adjusted *P*<0.01)^14^. This comparison led to the discovery of 564 CUGs and 250 CDGs that we considered as exhibiting ‘core tumor-specific’ expression pattern (**Fig. 3b-3c**). With respect to CUGs (**Fig. 3b**), 392 genes were low in the stroma (i.e., high in epithelium), but were not low in normal pancreas. This subset contained several genes described as by Moffitt et al. as tumor-specific, including *KRT7, AREG, S100A2, LYZ, TFF1, AGR2* and *CTSE*. This suggests that those genes though low in the stroma, are more expressed in other normal tissue types compared to the pancreas making them less ideal as drug targets. The other shown subset of genes (n=769) consisted of CUGs that are highly expressed in the stroma (**Supplementary Table 4a**).

We performed pathway analysis with the 564 ‘core tumor-specific’ CUGs and identified ‘cell cycle/division’, ‘glycolysis’, ‘p53’ and ‘progesterone-mediated oocyte maturation’ (consisting of *CCNB1, CDK1, HSP90AA1, MAD2L1, CCNB2, BUB1, CCNA2, CDK, CDC25B* genes), among pathways lowly expressed in normal pancreas and reprogrammed towards high expression in neoplasia (**Fig. 3d**). In contrast, the 250 CDGs revealed that normal pancreas expresses high level of genes in ‘protein processing in endoplasmic reticulum’ (e.g., *UBE2D4, ATF4, HERPUD1, MAN1A2, UBE2J1*), ‘FoxO signaling pathway’ (*IGF1R, SGK1, GABARAPL1, CCND2, NLK, FBXO25, BNIP3, FOXO3, GABARAP*) and ‘transcriptional misregulation in cancer’ (*IGF1R, NUPR1, CCND2, PDGFA, RXRA, PBX1, AFF1, PBX3, MEIS1*). Further, GO analysis showed that tumor-specific CUGs are associated with cytoplasmic, membrane and nuclear/nucleoplasmic activities (**Fig. 3e**), whereas tumor-specific CDGs are associated with endoplasmic reticulum (ER), golgi-apparatus and exosomes (**Fig. 3f**). Direct comparison of tumor to stroma using LCM data showed many CUGs and CDGs that are high and low, respectively, in the epithelial cells (**Fig. 3g-h**). Specifically, majority of the CUGs (48%, *n=956*) showed high expression in the epithelium compared to 21% that turned out to be elevated in the stroma. Similarly, although by a narrower margin, the majority of CDGs (18%, *n*=347) showed low expression in the epithelium compared to 11% in the stromal cells (**Supplementary Table 4a**). We recently published single cell RNA sequencing (scRNA seq) profile of human PDAC^16^. In this scRNA seq dataset, we called the epithelial population based on the expression of *KRT19* (a CUG) and also identified immune cell populations. However, while we confirmed the tumor-specific expression of several CUGs and found that some others also showed expression in immune cells (**Fig. 3i-j, Extended Data Fig. 3d-e**). Integrative analyses of these data therefore more accurately reveal tumor-specific and non-specific consistent genes that could guide and broaden the translational prospects of targeting specific or heterogeneously expressed signatures.

A major interest in cancer research is to determine gene alterations in tumors that may follow the same expression pattern in blood, and hence serve as potential biomarkers. We sought to determine the CUGs and CDGs that show high or low expression pattern in the peripheral blood mononuclear cells (PBMCs) of PDAC patients relative to healthy controls. To this end, we first determined the consistency of genes in microarray datasets of PBMC samples since that to our knowledge has not been previously reported. We observed that unlike in tumors, the blood datasets had considerably less differentially expressed genes, which led us to lower our selection criteria to *P*<0.05. Analysis of three datasets revealed a total of 1,592 reproducibly changed genes in at least 2 datasets (**Fig. 3k, Supplementary Table 4b, Note 2**). The pathway and GO profile of PBMC data were also remarkably different from tumor profile and were specifically oriented towards platelet biology and innate immune response for upregulated genes, and T-cell function, signaling pathways (e.g., ‘TGFB’ and ‘NFkB’) and cell cycle for downregulated genes (**Supplementary Table 4c**, **Fig. 3l**). Nevertheless, we identified 138 CUGs and 97 CDGs (including glycolysis genes, *MK167, MUC1/MUC4, TGFBR3, EIF3F/G/L*) that retained their tumor expression pattern in blood (**Fig. 3m-n**). These analyses identify consistent genes that display tumor-specific or microenvironment cell expression and represent probable biomarkers based of detectable expression in blood and tumor samples.

### The consistent genes correlate with tumor subtypes and aggressive features

We queried all tumor-specific or microenvironmental cell specific genes against classical and basal-like PDAC subtype classification as well as clinicopathological variables including proliferation, metastasis, mutation, tumor grade and patient survival outcome (**Extended Data Fig. 4a, Supplementary Note 2**). To determine the consistent genes associated with the basal-like and classical subtypes, we used 50 gene signatures of these subtypes published by Moffitt et al. ^10^ to stratify three patient cohorts (TCGA, Moffitt, and Puleo). Of note, ~70% of those 50 signature genes are among our consistent genes (**Extended Data Fig. 4b**). We found that in at least 2 cohorts, 2,101 and 1,779 genes (*P*<0.05) emerged as high in basal-like or classical tumor subtypes (**Supplementary Table 5**). We established that most of the overlapping CUGs (557 of 809, i.e., 69 %) were basal-like signatures and that 78% of CDGs (i.e., 273 of 350 overlapping hits) were classical subtype signatures (**Fig. 4a**). However, 1,503 of the total 3,938 CUGs/CDGs were not changed (*P*>0.05) between basal-like or classical subtypes in any of the three cohorts **(Extended Data Fig. 4c**). This suggests that there is at least one other subtype of PDAC that is non-basal-like non-classical or share features of both. By our estimation, this subtype represents ~38% tumor samples based on the number of genes. This subtype notably expresses high *TMPRSS4, SLC6A14, IFI27, CST1, STYK1 (CUGs*) and low *TMED6, PNLIPRP1, SERPINI2, ALB, AOX1* (CDGs), which are topmost consistent genes that neither mapped to basal-like nor classical subtypes (**Supplementary Table 5**). Of note, at the scRNA sequencing level, we hardly found a clear distinguishing pattern between the various subtypes (**Extended Data Fig. 4d**).

**Figure 4.**
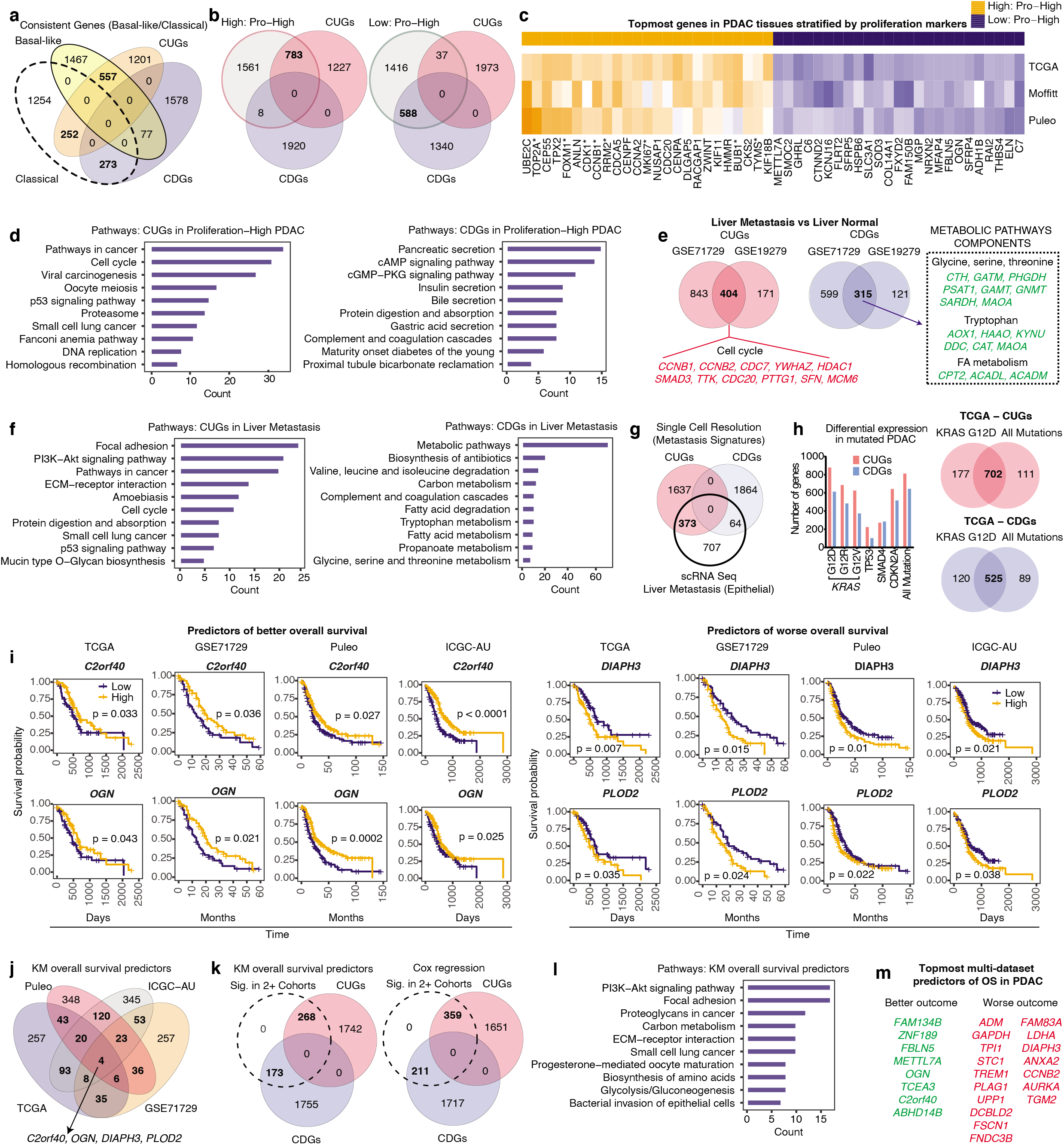
The consistent genes correlate with features of PDAC aggressiveness and poor prognosis. **a,** Venn diagram showing the number of core genes (in bold) overlapping between CUGs and CDGs in basal-like and classical subtypes of PDAC. The basal-like and classical subtype genes included were those high in basal-like relative to classical and vice versa in at least 2 of TCGA (*n*=31 basal-like vs 31 classical), GSE71729 (*n*=27 basal-like vs 27 classical) and Puleo (*n*=64 basal-like vs 64 classical tumor samples) datasets (*P*<0.05). **b**, Venn diagram showing the number of CUGs or CDGs overlapping with differentially expressed genes in proliferation (‘pro’)-high tumors. Genes were selected as altered in proliferation-high tumors if differentially expressed (*P*<0.05) relative to proliferation-low tumors in at least 2 of TCGA (*n*=64 pro-high vs 86 pro-low), GSE71729 (*n*=77 pro-high vs 68 pro-low) and Puleo et al. (*n*=99 pro-high vs 210 pro-low) tumors (*P*<0.05). **c**, Heatmap depicting the topmost genes consistently high in at least 2 of the 3 datasets stratified by proliferation markers. *proliferation markers used for the tumor stratification; included as positive controls. **d**, Pathway annotation of the 783 CUGs and 588 CDGs that overlapped with genes differentially expressed in proliferation-high vs low tumors in at least 2 datasets. **e**, Venn diagram showing CUGs and CDGs overlapping with genes in liver metastasis compared to normal liver tissues from GSE71729 and GSE19279 datasets (*P*<0.05). See Methods for sample sizes/types. **f**, Pathway annotation of the 404 CUGs and 315 CDGs overlapping in the liver metastasis compared to normal liver tissues from GSE71729 and GSE19279 datasets. **g**, Single cell RNA sequencing data showing the overlap between CUGs or CDGs and epithelial gene signature from liver metastasis samples (*n*=5 tumors tissues). **h**, Number of differentially expressed genes in tumors with mutation versus tumors with no alteration in the indicated genes in TCGA data. “All mutation” refers to tumors with mutation in any of *KRAS, TP53, SMAD4, CDKN2A* versus tumors with no recorded mutation. On the right, overlap of genes differentially expressed in KRAS G12D tumors versus in “All mutation”. **i,** Kaplan-Meier (KM) overall survival plots (log-rank test) of genes that predicted survival in the clinical cohorts analyzed. Tumor sample size, TCGA (*n*=146), Puleo et al (*n*=288); ICGC (*n*=267); GSE71729 (*n*=125). **j**, Venn diagram showing overlaps of KM OS predictors in TCGA, GSE71729, Puleo et al. and ICGC-AU cohorts. **k**, Venn diagram showing the number of CUGs and CDGs that predicted OS or hazard ratio (univariate Cox regression) in at least 2 out of TCGA, GSE71729, Puleo et al. and ICGC-AU cohorts. Sig – significant (*P*<0.05). **l**, Pathway annotation of the 441 genes that predicted OS in at least two datasets. **m**, Topmost genes that predicted overall survival in PDAC. These genes predicted OS in at least 3 of the 4 datasets mentioned above and were ranked based on the sum of their log (*P*-value) across the datasets. Log-transformation was applied only for *P*<0.05.

We also stratified the same three patient cohorts into high and low proliferation subsets using 26 known proliferation-associated markers. Interestingly, over 2,000 genes overlapped as differentially expressed between proliferation high/low in ≥ 2 cohorts, indicating the high reproducibility of these stratification results (**Fig. 4b, Supplementary Table 5)**. We found that a preponderance of CUGs (39%, *n*=783) were high in tumors expressing high proliferation markers whereas <2% (n=37) were high in proliferation-low tumors. In addition, the topmost genes in proliferation-high tumors were notably CUGs (**Fig. 4b-4c**). The proliferation-high CUGs were clearly enriched in ‘pathways in cancer’, ‘cell cycle/division’, ‘p53’ and proteasome, indicating the processes underlying tumor proliferation (**Fig. 4d**). Some CDGs, e.g., *PSAT1* in serine metabolism, *C9orf40, GPR3, PRPS2, SAMD1, IPO4, GIT1* and *DFFA* surprisingly emerged as high in proliferation-high tumors despite their low expression in tumor relative to non-tumor samples (see **Supplementary Note 3**). Otherwise, majority of CDGs (n=588, ~31%) were low in proliferation-high PDACs and these include genes such as superoxide dismutase 3 (*SOD3), OGN, ALDH1B, SLC3A1* and complement components (*C6/C7*) (**Fig. 4d**). These CDGs were strikingly associated with physiologic pancreatic activities, e.g., ‘gastric and insulin secretion’, ‘protein digestion/absorption’, altogether revealing for the first time the processes whose differential gene changes underlie PDAC proliferation across cohorts.

Metastasis accounts for >90% of cancer deaths and is a major feature of PDAC^17^. We analyzed three PDAC microarrays with metastasis samples and performed over 10 comparisons of metastatic versus primary PDAC or distant normal versus lymph nodes, peritoneal lung, or liver metastasis tumors (**Supplementary Table 6, Supplementary Note 3**). We found that several topmost CUGs in PDAC, e.g., *LAMB3, LGALS4, CEACAM6, TFF2* were also the top upregulated genes in metastasis tissues (**Extended Data Fig. 4e, Supplementary Table 6**). The liver showed the highest overlap with CUGs. Specifically, in the GSE71729 dataset, 91% of the CUGs differentially expressed in liver metastasis relative to normal liver were high. This is followed by the lymph node at 87% and lung metastasis 82%. Thus, >80% of CUGs are high in metastasis. Analysis of the overlapping genes between GSE71729 and GSE19279 revealed 404 highly expressed CUGs and 315 lowly expressed CDGs in liver metastasis (**Fig. 4e**). Pathway and GO analyses revealed that the ‘metastasis-CUGs’ are involved in ‘focal adhesion’, ‘extracellular matrix-receptor interaction’, ‘p53’, ‘cell cycle’, ‘proliferation’ and ‘mitochondrial matrix’ - with notable less involvement of glycolysis (**Fig. 4f**). On the other hand, the CDGs lowly expressed in liver metastasis were enriched for the downregulation of metabolic pathways, which was clearly different from the pancreatic physiology-associated profile noticed with the CDGs low in proliferation-high PDAC (**Fig. 4e-f**). Topmost CDGs in PDAC, e.g., *FCN2/FCN3, CXCL12, CTSG*, and lymphatic endothelial marker *LYVE1*, also showed a low expression across various metastatic samples (**Extended Data Fig. 4e, Supplementary Table 6**). We also found several CUGs and CDGs that strongly overlapped between liver and peritoneal metastasis samples (**Extended Data Fig. 4f**). Of note, the non-basal-like non-classical signatures at 8.9% (n=134) had more overlapping genes with metastasis than basal-like (115, n=5.5%) and classical subtypes (3.3%, n=58) (**Extended Data Fig. 4g**), further suggesting that this tumor subset could be more aggressive. At the singlecell resolution, we observed that >370 CUGs were highly expressed in the epithelial cell compartment of liver metastasis tumors (**Fig. 4g**). Altogether, these data reveal the most robust gene signatures associated with PDAC metastasis, many of which have never been reported in any PDAC context.

The most frequently mutated genes in PDAC are known (i.e., *KRAS, TP53, SMAD4* and *CDKN2A*), but their impact on tumor gene expression pattern is not well-defined. We analyzed how tumors with the mutated genes differ in CUG- and CDG expression using TCGA data, and for each gene we separately compared tumors with mutation to those without. We found that *KRAS*-mutated tumors expressed high *KRAS, SMAD4-mutated* tumors expressed low *SMAD4*, but there was no change in *TP53* or *CDKN2A* levels for tumors harboring mutations in *TP53* or *CDKN2A*, respectively. Also noteworthy, regardless of the gene mutation considered, *SMAD4* expression level was low (**Supplementary Table 7a**). *KRAS* mutation, mainly G12D, exerted the most profound impact on the CUGs and CDGs, followed by mutations in *CDKN2A* and *SMAD4* (adjusted *P*<0.05, **Fig. 4h, Supplementary Note 3**). *KRAS* G12D alone almost exclusively accounted for all the differentially expressed CUGs and CDGs in mutated tumors (**Fig. 4h**), and we identified 418 overlapping CUGs and 286 CDGs altered in *KRAS* G12D, G12R and G12V tumors (**Extended Data Fig. 4h, Supplementary Table 7b)**. *TP53* mutations had the least impact probably because, unlike *KRAS* mutations which could easily be grouped into G12D, G12R and G12V, almost every 2-3 samples had a different *TP53* mutation indicating extensive heterogeneity. Pathway analysis of CUGs in *KRAS* G12D-mutated tissues identified ‘cell cycle’, ‘p53’, ‘glycolysis’ (**Extended Data Fig. 4i)** among other alterations that were repeatedly observed in the proliferation and metastasis comparisons. Thus, we conclude that of the gene mutations, KRAS mutation is the main orchestrator of the consistent gene expression changes observed in PDAC.

We used TCGA and Puleo cohorts to further determine CUGs and CDGs that specifically distinguish poorly differentiated (Grade III) from moderately differentiated (Grade II) tumors, considering that such genes could complement histological grading and fine-tune diagnostic precision. We surprisingly found few overlapping genes (178 upregulated and 138 downregulated genes, *P*<0.05) between Grade III relative to Grade II tumors in both datasets. Nevertheless, of these genes, 58 upregulated genes in Grade III tumors were CUGs, including *ZBED2, S100A2, KRT6A, IL20RB, MET*, glycolysis genes (*PFKP, LDHA*), uridine phosphorylase 1 (*UPP1*) (**Supplementary Table 8**), suggesting indicators of poorly differentiated tumor grade. Several CUGs also showed specificity for Grade III based on receiver operating characteristics (ROC) area under the curve (AUC) analysis (**Supplementary Table 8**). Only 10 CDGs, notably *NDRG2, DCDC2, PDX1, MYOM1, ZFPM1, ACVR1B*, emerged in the Grade III vs II comparison (**Supplementary Table 8**). The Puleo cohort contained many Grade I tumors (*n*=117 samples), which enabled us to further predict CUGs or CDGs associated with progression from well to poorly differentiated tumor stages and also compare Grade II vs I tumors. We identified 10 CUGs (including *S100A2, HMGA1/2, WNT7B, RDX*) and 21 CDGs, including *PLA2G10, CREB3L1, ATP10B, TOX3, SLC37A1* and *GATA6* from ROC AUC analysis. Pathway analysis of Grade II vs I tumor signatures showed ‘focal adhesion’, ‘Wnt’ and ‘PI3K-Akt’ among processes in involved in the initial stages of tumor progression (**Extended Data Fig. 4j**). In an independent cohort of pre-malignant PDAC samples ^18^, we also observed differential expression of CUGs and CDGs (**Extended Data Fig. 4k, Supplementary Table 9**), which suggest that several of these alterations occur early in the tumor progression course.

One common utility of gene expression data is for predicting patient overall survival (OS) outcome and hazard ratios. To determine which CUGs and CDGs best predict OS or risk of mortality, we performed Kaplan-Meier OS and univariate Cox regression analyses, respectively, for all genes in TCGA, Moffitt, Puleo and ICGC-AU cohorts (**Supplementary Table 10a, Supplementary Note 3**). We identified four consistent genes, namely *OGN*, *C2orf40* (CDGs) and *PLOD2*, *DIAPH3* (CUGs) that predicted OS in all four datasets. Indeed, 1,648, 441 or 61 CUGs/CDGs predicted OS in at least 1, 2 or 3 cohorts, respectively (*P*<0.05) (**Fig. 4i-j**). Further, the number of CUGs and CDGs that predicted OS in at least two cohorts largely mirrored the result of univariate Cox proportional hazards regression (**Fig. 4k**), and in an independent dataset of ‘bad’ versus ‘good’ prognosis tumors several CUGs/CDGs also emerged (**Extended Data Fig 4l, Supplementary Table 10b**). Of note, ‘PI3K-Akt’, ‘focal adhesion’, ‘carbon metabolism’ and ‘glycolysis’ were the topmost pathways associated with the CUGs that predicted OS (**Fig. 4l**). Ultimately, we identified 25-30 predictors of OS in PDAC as the most robust, based on significant prediction in ≥ 3 cohorts. These predictors in multi-datasets included *OGN, PLOD2, DIAPH3, C2orf40* (shown in **Fig. 4i**), *ALDOA, GAPDH, PKM2, LDHA* (in glycolysis), *ADM*, *UPP1, ZNF189, DCBLD2, PLAG1* among others (**Fig. 4m**). Of note, for some genes a high expression predicted better outcome in a cohort, but worse outcome another indicating potential context-dependent caveats often not captured when using a single dataset (**Supplementary Note 3**). Taken together, we present the most extensive PDAC-centric crossplatform analysis revealing gene expression pattern associated with clinical variables in patients.

### High priority consistent gene targets predict epigenetic inhibitors for PDAC therapy

To determine high priority therapeutic targets among the consistent genes, we analyzed the CRISPR/Cas9 knockout screen data from Project Achilles^19^, GECKO and Behan et al^20,21^ studies, and also analyzed the Project Drive^22^ shRNA knockdown screen data (**Fig. 5a**). The original publications used these RNA interference (RNAi) screens to determine the essential genes for growth/survival across human cancer cells/types. We first checked the essential gene results from these studies considering that they did not directly focus on PDAC. We found that of the genes identified by the Project Achilles as essential across > 700 cell lines, 245 overlapped with our CUGs and 137 were CDGs. Also, over 400 CUGs overlapped with the cancer fitness genes defined by Behan et al. Further, 972 CUGs were among genes included in the Project Drive shRNA screen, which originally focused on ~8000 genes that the authors predicted to be essential in most cancers (**Supplementary Table S11a**). These observations reinforced our confidence that several consistent genes we identified are essential for the viability of PDAC and ostensibly in multiple other cancer cell lines and types. In addition, the possibility of querying RNAi of between eight to ~24 PDAC cell lines across four independent studies offered unprecedented statistical strength to our analyses.

**Figure 5.**
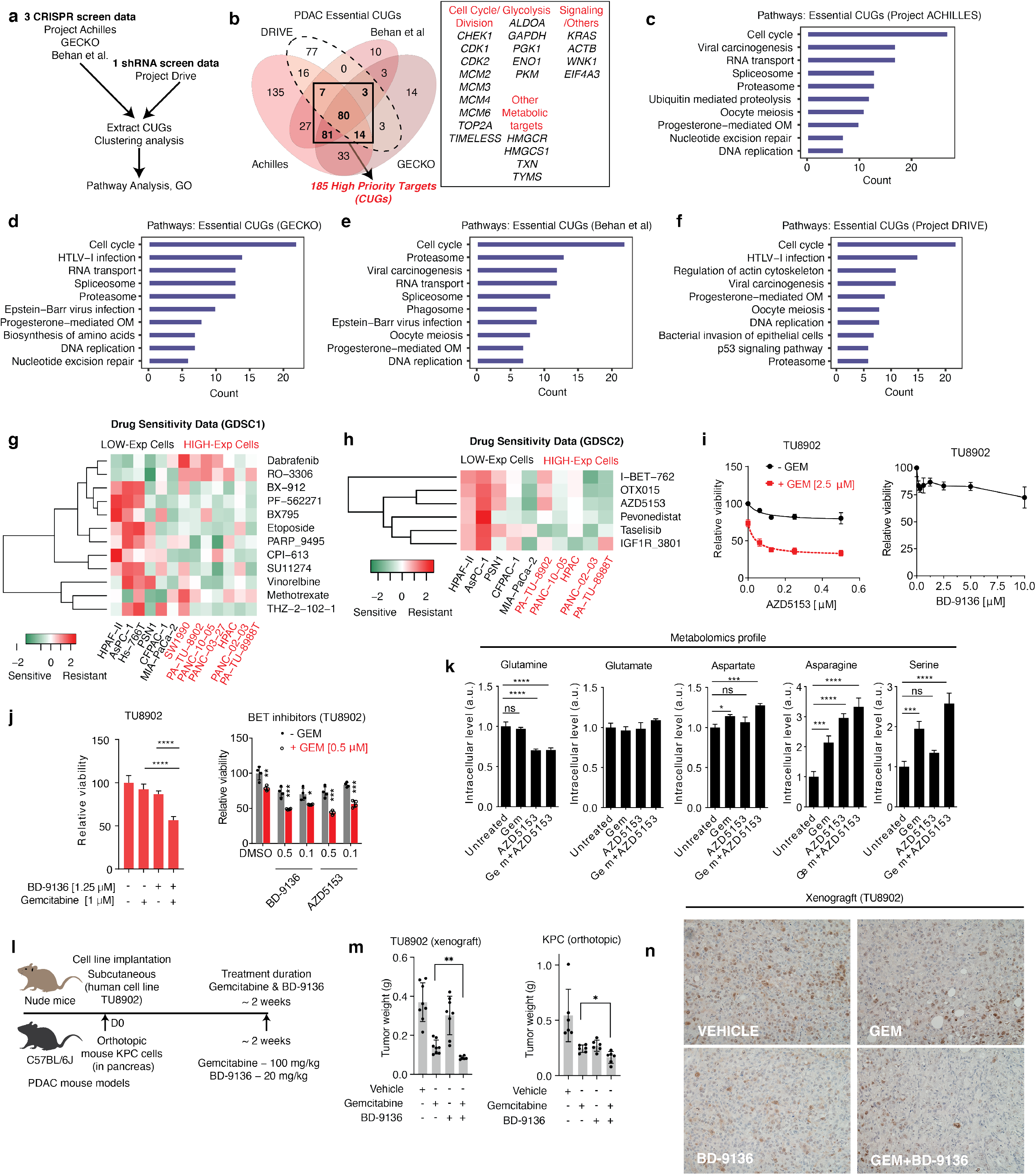
The high priority therapeutic targets predict epigenetic inhibitors for PDAC. **a**, Schematic overview of the workflow for the RNA interference (RNAi) screen data analysis. **b**, Venn diagram showing the overlapping essential genes in PDAC cell lines as derived from the Project Drive, Project Achilles, Behan et al. and GECKO screen data. In bold are the number of CUGs, in total 185, that overlapped as essential for survival in at least 3 of the 4 RNAi screen data. On the right are selected components among the 185 high priority targets. **c-f**, Pathway annotation of genes that emerged as essential for PDAC survival/growth in Project Achilles (*n*=394 CUGs), GECKO (*n*=231 CUGs), Behan (*n*=211 CUGs) and Project Drive (*n*=200 CUGs). **g-h**, Heatmaps showing the sensitivity of PDAC cell lines expressing high priority genes to various compounds tested in the cancer drug response project. **i**, Viability assay of PDAC cell lines treated with BETi AZD5153 alone or in combination with gemcitabine for 72h. On the right, viability assay of TU8902 cells treated with BETi BD-9136 at high concentration. **i**, Viability assay of TU8902 cell line treated with gemcitabine, BETi BD-9136 or both (48 h). Treatments were in quadruplicates; representative of >2 independent experiments. On the right, treatment with BD-9136 or AZD5153 alone or in combination with gemcitabine for 48 h. **l**, Metabolomics profiling showing amino acids (intracellular abundance) disrupted by the inhibitors after 24 h treatment. Data normalized to untreated control; a.u.- arbitrary unit; experiment was in triplicates. Schematics of the mouse experiment. Mice were treated with gemcitabine at 100 mg/kg body weight 2x/week and with BD-9136 at 20 mg/kg body weight 5 times/week. **m**, Tumor volume of subcutaneously implanted TU8902 cell lines (*n*=4 mice per group, injected on both flanks, *n*=8 tumors per arm) and KPC (7940b cell) orthotopic tumors (*n*=6 mice per group). Figures 5j and l, * *P*<0.05, ** *P*<0.01, **** *P*<0.0001. **n**, Representative micrograph (20X) of Ki-67 staining of the xenograft tumor samples derived from TU8902 cell lines. Figures 5j, k and m – statistics by analysis of variance (ANOVA).

To determine the most probable therapeutic targets in PDAC, we focused on the CUGs since it is more logical to hypothesize that suppressing their expression will suppress tumor growth. Clustering analysis of Achilles and Project Drive (each containing 24 PDAC cell lines) identified 394 and 200 CUGs, respectively (**Supplementary Table 11a**). In the GECKO dataset (*n*=8 PDAC cell lines) we identified 231 CUGs of high essentiality and found 211 in the Behan et al dataset (*n*=20 PDAC cell lines). On average, we identified ~206 essential CUGs across the four RNAi screens, of which 185 CUGs overlapped as essential in at least 3 datasets (**Fig. 5b**). These most reproducible CUGs (herein called “high priority therapeutic targets”) included glycolysis genes (*ALDOA, PKM, GAPDH*), cell cycle/division genes (*BUB1, CDK1, CDK2, MCMs*), thioredoxin (*TXN), KRAS*, and *ACTB* (actin beta) involved in cell motility/membrane integrity. Pathway analysis revealed a remarkable similarity between the identified priority targets per dataset (**Fig. 5c-f**). Specifically, ‘cell cycle/division’, ‘proteasome’, ‘spliceosome’ and ‘progesterone-mediate oocyte maturation’ were the most consistent pathways depicted by the high priority targets (**Supplementary Table 11b**). Others included ‘p53’, ‘regulation of actin skeleton’ (**Fig. 5d, 5f**) as well as ‘biosynthesis of amino acids’ (**Fig. 5d**), which actually contained mainly glycolysis genes. GO analysis identified the biological process ‘cell cycle’, molecular function ‘protein and ATP binding’, and cellular component ‘nucleoplasm,’ ‘cytosol’ and ‘cytoplasm’ as key ontologies linked to priority target CUGs (**Supplementary Table 11b**).

To predict therapeutic agents that could be effective against PDAC, we used the 185 high priority targets to stratify PDAC cell lines in the cancer cell line encyclopedia (CCLE) and GSE57083 datasets into two groups (**Extended Data 5a**). Representative PDAC cell lines that best showed a higher expression of the priority genes included PA-TU-8902, PANC1, PANC-03-27 (i.e., the ‘high-expressing’) relative to AsPC-1, CFPAC-1, and HPAF-11 (low expressing). We analyzed the sensitivity of these cells to therapies using data from the Genomics of Drug Sensitivity in Cancer (GDSC) project^23^ and identified several drugs that were effective against cell lines expressing high priority targets (**Fig. 5g**). Interestingly, bromodomain and extra-terminal motif (BET) protein inhibitors I-BET-762, OTX015, and AZD5153 emerged as effective in most high-expressing cell lines with the notable exception of PA-TU-8902 indicating a potentially resistant cell line (**Fig. 5h**). BETi act by repressing transcription and cell cycle^24,25^, which are core alterations associated with CUGs in PDAC. Given that gemcitabine, a major chemotherapy agent used for PDAC therapy, acts by suppressing nucleotide biosynthesis and cell cycle, we tested the combination of gemcitabine and BETi in TU8902 and other cell lines. We found that the cell lines were not strikingly sensitive in their response to BETi alone. However, BETi synergized with gemcitabine to suppress cell viability across all the tested cell lines (**Fig. 5i-j, Extended Data Fig. 5b**). Metabolomics revealed a strongly disrupted amino acid metabolism upon combination treatment, with the notable accumulation of serine (**Fig. 5k, Extended Data Fig. 5c-d**), which we previously observed is an indicator of nutrient deprivation and suppressed proliferation^26^. In both a human cell line subcutaneous xenograft model with TU8902 cell line, and in immunocompetent orthotopic mouse model with oncogenic *Kras; Trp53* R172H/+ cells (KPC), the combination of gemcitabine and BETi BD-9136 synergistically suppressed tumor growth (**Fig. 5l-m, Extended Data Fig. 5e**). This was accompanied by a suppressed proliferation as indicated by Ki-67 staining (**Fig. 5n**). We conclude that the high priority genes are themselves useful targets for therapy and cell line stratification by their expression predict that targeting epigenetics could be an effective combinatorial therapeutic option against pancreatic tumors.

## Discussion

Genomic profiling has paved way for unprecedented discoveries in many cancers^27,28^. For the first time, we have extensively leveraged on multiple published datasets to define the most consistently expressed genes in human PDAC – providing a refined set of genes for future research. These genes are unambiguous in their expression pattern and can be expected to be significantly changed in any comparison of PDAC tissues to non-tumoral pancreas. Over 50% of the 3,938 identified consistent genes are not functionally characterized in PDAC revealing untapped opportunities for future new discoveries. Among the consistent genes, those previously studied were often proposed as therapeutic targets, biomarkers in PDAC or were shown to exhibit the same pattern as described in our work, providing additional confidence in the potential of the uncharacterized genes/drug targets. Further, several genes previously identified as tumor/stroma specific or signatures of basal-like and classical PDAC subtypes^10,14,29^, also emerged in our analysis of multiple datasets. These observations reinforce confidence in the strength of our findings and align with our conclusion that these consistent genes are plausibly important in PDAC. We show that the identified consistent genes correlated with several clinicopathological indices, are detectable in blood, and contain high-priority essential therapeutic targets in PDAC, which altogether broaden the utility of the genes in basic and translational research.

Certain key observations are noteworthy. We found several known genes (e.g., *HIF1A, SLC16A3, EPCAM* and *SOX9*) that retained a consistent expression pattern but did not meet our inclusion criteria. Issues of gene probe identification also may have caused some genes, e.g., *KRT17* and *INHBA*, to fail our consistency test. Accordingly, we believe our estimate of ~4000 consistent genes in PDAC, though numerous at face value, is conservative and underlies the molecular complexity of this lethal disease. We recommend the use of analytical tools for pathway enrichment and gene ontology^30^ when the goal is to determine individual overrepresented pathways that may be investigated in experimental contexts. We also note that not all consistent genes followed the same expression pattern across comparisons. For example, several CUGs were indistinctly expressed in normal pancreas compared to other tissues and some were even higher in the normal pancreas. This was also notable with glycolysis genes, which have been traditionally considered high in most cancer types. For example, in single cell RNA sequencing data, we found that several core glycolysis genes *ALDOA, GAPDH, ENO1*, and *PKM2* are highly expressed in both the tumor and immune cell compartments. We believe such CUGs should not be considered tumor specific, but that tumor nonspecificity does not render them less important. Indeed, correlation with proliferation, metastasis and other clinical parameters also showed that in those comparisons some tumor non-specific consistent genes were nevertheless significantly changed – with glycolysis genes strongly appearing as the best candidates in most comparisons. Therefore, our identification of tumor-specific and non-specific genes could help facilitate hypothesis generation for studies targeting or exploring biomarkers of both compartments.

Next, by stratifying tumors using signatures of basal-like and classical subtypes previously published by Moffitt et al^10^, we establish that PDAC does not easily bin into two subtypes. Our data indicate that there exist at least one other subtype that is neither basal-like nor classical or at best combines features of both subtypes. We propose calling those subsets ‘non-basal-like non-classical’ and we observe that their core signatures include the high expression of *TMPRSS4, SLC6A14, IFI27, CST1* and *STYK1*. Of note, *IFI27*, a novel target in PDAC that was a strong marker of epithelial cell compartment in our analysis has been implicated in metastasis in two recent studies^31,32^ and we find that the non-basal-like non classical signatures more overlap with liver metastasis signatures, suggesting that this subtype may be the most aggressive. It is also noteworthy that basal-like and classical subtype signatures are hardly distinguishable in their expression pattern at the single cell level. Specifically, several signatures of both subtypes are either strongly expressed in the epithelial cell compartment or showed minimal expression. Thus, the mechanistic roles of PDAC subtype signatures are likely context dependent. In the context of OS, some genes predicted better outcome in one cohort and worse outcome in another. This pattern might reflect differences in patients’ clinical course or tumor subtypes and suggest that observations of survival prediction should not be generalized without testing in independent cohorts. Our findings also demonstrate that although some pathways, e.g., cell cycle and glycolysis, consistently ranked top, alterations in PDAC are hardly attributable to just one pathway. For example, although ‘focal adhesion’, ‘p53’, ‘ECM-receptor alterations’ and ‘pathway in cancer’, are the most prominently upregulated pathways from CUGs, other enriched pathways emerged in subsequent comparisons such as in metastasis, survival and notably PBMCs samples.

Based on data from normal pancreas, proliferation-high tumors, RNAi, and mutational landscapes, we propose that the high expression of glycolysis- and cell cycle/division pathway components are at least two persistent alterations governing the origin and progression of PDAC. This notion is consistent with a prior study, which found that genomic rearrangements and alterations in cell cycle likely occur at the early stage of PDAC^17^. Glycolytic and cell cycle/proliferation profile of normal pancreas are broadly low but are high in almost every tumor comparison we performed. The notion that glycolysis drives cancer is the foundation of the century-old Warburg effect, which recently was substantiated to promote both tumor and immune cells in the microenvironment^33^. Despite the long-standing appreciation of glycolysis, the molecular details remain incompletely defined. For example, the role of specific glycolytic enzymes or their isoforms e.g., *HKs, PFKP* and ENO2, remain unclear, and the therapeutic prospects of inhibiting these individually and in combination are largely unknown. Similar to glycolysis, despite the prevailing acceptance of cell cycle as a major cancer hallmark, most cell cycle genes we identified have never been studied in PDAC. We also identified *KRAS* among the priority genes. Specific inhibitors for KRAS for human studies are lacking and targeting the pathway component was ineffective in prior clinical trial ^34^. However, the continual emergence of *KRAS* in PDAC studies, coupled with experimental evidence of its role in tumorigenesis in mice^6,7^, suggest that efforts should continue to explore new ways to target *KRAS*. Other processes such as ‘proteasomes’ and ‘spliceosomes’ contained consistent genes in PDAC that showed essentiality across high-throughput RNAi screens^19,20,22^. These genes are also low in normal pancreas and hence are among the most attractive unexplored PDAC therapeutic target.

Lastly, we have demonstrated the utility of these consistent genes in predicting therapeutic options in PDAC. We predict that inhibitors of epigenetic process, namely inhibitors of bromodomain containing proteins, could be effective in PDAC. This observation, which we confirmed in two different *in vivo* models, will require further exploration. Besides predicting therapies, these genes are potentially direct therapeutic targets. In conclusion, we have performed the most comprehensive multi-dimensional analysis of gene expression pattern in PDAC to date and have unraveled >2000 consistent genes not yet functionally characterized or associated with this disease. Our data is a novel knowledge base and resource for high confidence basic, translational, and clinical studies. It is expected to revolutionize future understanding of true drivers of PDAC and facilitate the application of such insight in improving patient care.

## Methods

### Identification of the consistent genes

The following five human PDAC microarray datasets were used for the initial identification of the consistent genes: GSE71729 (46 normal pancreas, 145 tumor tissues), GSE62452 (61 non-tumoral vs 69 tumor tissues), GSE28735 (45 non-tumoral vs 45 tumor tissues), GSE16515 (16 normal vs 36 tumor tissues) and GSE15471 (36 non-tumoral vs 36 tumor tissues). These datasets were obtained from the National Center for Biotechnology Information gene expression omnibus (NCBI GEO) (https://www.ncbi.nlm.nih.gov/geo/). All datasets were analyzed separately. For each dataset, the differentially expressed genes in pancreatic tumors compared to non-tumoral control samples were determined using *limma* package (v 3.38.3) in R software (v 3.5.2) after quantile normalization. Where there are multiple probes identifying a particular gene, we obtained the average for that gene prior to analysis. We used adjusted *P*<0.05 for selecting differentially expressed genes per dataset. Genes that met the adjusted p-value cut off, and were upregulated in at least four of the five datasets, were included as consistently upregulated genes (CUGs). Similarly, genes that showed low expression in at least four datasets were included as consistently downregulated genes (CDGs). Subsequently, we generated a matrix file containing the significantly changed consistent genes for each of the five datasets (*n*=2010 CUGs, and *n*=1928 CDGs in total). For some gene symbols and alias (e.g. *PKM2*), conversion to current human gene nomenclature was necessary for cross-dataset comparison. For that we used either the https://www.biotools.fr/ portal or *limma* built-in function for alias conversion. The consistent genes were ranked using their average ‘expression score’ (i.e., the product of logFC and -log *P*-value) across the five datasets. Besides the list of CUGs and CDGs (expressed in at least four datasets), we extracted genes that showed consistent expression in at least three datasets, provided they are unchanged in the remaining two datasets (*n*=2,722, adjusted *P*<0.05, see **Supplementary Table 2**). These genes were only used in completing metabolic pathway drawings in cases where they are critical components of the pathway but did not meet selection cut-off of the CUGs and CDGs list described in this work. Where applicable, those genes were marked with asterisks.

### Cross-validation of consistent genes in independent tumor and blood datasets

For cross-validation of CUGs and CDGs in independent patients cohorts, we included four additional microarray datasets, namely, GSE32676 (7 non-malignant pancreas vs 42 tumor tissues, *P*<0.05), GSE62165 (13 control samples vs 118 tumors, adjusted *P*<0.01), and GSE19279 (compared 3 normal vs 4 tumors, *P*<0.05) and GSE19650 – a laser capture microdissection data, 7 normal pancreas vs pre-malignant tissues of invasive cancer originating in intraductal papillary-mucinous neoplasm (IPMN, *n*=3), intraductal papillary-mucinous adenoma (IPMA, *n*=6), and intraductal papillary-mucinous carcinoma (IPMC, *n*=6), *P*<0.05). For analysis of differential gene expression in peripheral blood mononuclear cells (PBMCs) and comparison with the tissue data, we used three datasets: GSE15932 (8 healthy control vs 8 samples from non-diabetic PDAC patients), GSE49641 (18 healthy controls vs 18 PDAC patients), and GSE74629 (14 healthy control vs 22 non-diabetic PDAC patients). These datasets were processed and statistically compared following same methods as for tumors. Of note, some of these datasets have additional sample groups that are not necessarily tumors vs non-tumors. Where those additional samples were analyzed (e.g., metastasis vs distant normal tissues in GSE71729, we followed the analysis steps used for tumor vs non-tumors). Additional microarray datasets included were E-MTAB-6134 (309 tumors) obtained from ArrayExpress (https://www.ebi.ac.uk/arrayexpress/) as well as the International Cancer Genome Consortium – Australia cohort (ICGC-AU, 269 tumors). These two data were mainly used for tumor stratification, clustering and analysis of correlation with clinical parameters. Accession numbers of these datasets are indicated in **Extended Data Fig. 1a**.

### RNA Sequencing data

The patients RNA seq data we used were The Cancer Genome Atlas (TCGA) PDAC data (150 tumors) downloaded from cBioPortal (https://www.cbioportal.org/) and a laser microdissection (LCM) data GSE93326 (65 epithelium vs 65 stroma) downloaded from NCBI GEO. We retained only genes with an expression value ≥1 in at least half of the samples contained per dataset. To determine differential gene expression, we analyzed TCGA data with *limma* package after log2 transformation, whereas the raw count values for GSE93326 were analyzed with *DESEq2* package (v 1.22.2) in R. Unless otherwise indicated, adjusted *P*<0.05 was used as cut off for differential gene expression. We also included the normal tissue RNA seq from the Human Protein Atlas (HPA, 37 tissues including the pancreas) and the Genotype-Tissue Expression (GTEx, 36 tissues including the pancreas). The HPA and GTEx data were both downloaded from the HPA archive version 19 (https://www.proteinatlas.org/) and used as described below.

### Single-cell RNA sequencing analysis

The determination of the consistent gene expression at the single-cell level was done using our recently published human sample data from PDAC to liver metastasis (*n* = 5)^35^ or adjacent normal (*n*=3) versus PDAC tumor samples (*n*=16)^16^. For the metastasis samples, we first derived genes differentially expressed in the tumor epithelial compartment relative to other microenvironmental cell population. These genes were subsequently overlapped with the CUGs and CDGs to extract the common genes.

### Determination of normal pancreas expression of the tumor-derived consistent genes

To determine lowly expressed CUGs or high GDGs in normal pancreas, we used the HPA and GTEx data (each contained ≥ 97% of the consistent genes). We extracted the CUGs/CDGs from each dataset and calculated their median expression values for all other tissues except the pancreas. Genes that their expression in the pancreas was lower than the median for all other tissues were considered lowly expressed in the pancreas, and *vice versa*. Genes that showed low- or high expression in both the HPA and GTEx datasets were selected as consistently low or highly expressed genes in the pancreas. We also analyzed normal tissues in the GSE71729 dataset. Specifically, we compare the combined gene expression of normal liver (*n*=27), lung (*n*=19), lymph node (*n*=10) and spleen (*n*=11) to that of normal pancreas (*n*=46, *P*<0.01). Genes with the same expression pattern in GSE71729 as in HPA/GTEx were selected as highly consistent in normal tissues.

### PDAC stratification by proliferation, basal-like and classical subtypes

For the proliferation high-versus low tumor identification and comparison, we first generated tumor sample data subset containing the expression values of 26 known proliferation genes (**Extended Data Table 2**) from each of TCGA (*n*=150), GSE71729 (*n*=145), and Puleo (*n*=309) datasets. On each subset, we separately applied unsupervised hierarchical clustering and partitioned the tumor samples into proliferation high- and low clusters. Sample numbers were as follows: TCGA – 64 proliferation-high vs 86 low; GSE71729 – 77 proliferation-high vs 68 low; and Puleo – 99 proliferation-high vs 210 low. For generating the basal-like and classical subtype subsets, we used the 25 basal-like and 25 classical gene signatures published by Moffitt et al^10^. These genes were applied to rank each of TCGA, GSE71729 (Moffitt) and Puleo datasets into the two subsets. We then used *limma* package to derive the differentially expressed genes between compared groups. CUGs that emerged as significantly upregulated in at least two of the three datasets (*P*<0.05) were considered high in proliferation-high tumors and *vice versa*. The same rule was applied for CDGs.

### Correlation of consistent genes with metastasis

We used three bulk tumor datasets, namely, GSE19279, GSE42952, and GSE71729 to determine the consistent genes correlated with metastasis. With GSE19279 our analysis was liver metastasis samples (*n*=5) vs liver normal (*n*=3) or primary PDAC (*n*=4). With GSE42952, we compared tumor samples of liver metastasis (n=7) vs tumor samples (*n*=6) from patients that had ‘good’ prognosis (i.e., longer survival time, *n*=6) or ‘bad’ prognosis (i.e., shorter survival time, *n*=6). On the same dataset, the ‘good’ or ‘bad’ prognosis samples were also separately compared to peritoneal metastasis samples (*n*=4). With GSE71729, the following comparisons were performed: liver metastasis (*n*=25) vs distant site normal liver tissue samples (*n*=27); lung metastasis (*n*=8) versus normal lung samples (*n*=19) and lymph node metastasis (*n*=9) vs normal lymph node samples (*n*=10). In addition, we compared each of the mentioned metastasis tissue samples to primary PDAC (*n*=145). The CUGs and CDGs differentially changed in the respective comparisons (*P*<0.05) were identified and are presented in **Supplementary Table 6**.

### Differential expression of CUGs/CDGs based on frequently mutated genes

For the impact of frequently mutated genes in PDAC (i.e., *KRAS, TP53, SMAD4* and *CDKN2A*), we extracted the mutation profiles for each tumor sample in TCGA data as obtained from cBioPortal. For KRAS, we analyzed for G12D, G12R and G12V since these had considerable sample sizes. Specifically, we performed the following comparisons: tumors with G12D mutation (*n*=42), G12V mutation (*n*=26) and G12R mutation (*n*=22) each separately compared to no alteration in KRAS (*n*=43). For TP53, there were 80 indicated unique mutations of which the most represented (i.e., R175H and R248W) had four samples each. Therefore, we compared all tumors with any TP53 mutation (*n*=100) to those with no alteration (*n*=50). The same approach was taken for SMAD4 (55 mutation vs 95 no alteration) and CDKN2A (80 mutation vs 70 no alteration). Furthermore, we compared tumors with no alteration in any of these four genes (*n*=23) to tumors with a mutation in all four (*n*=24). Determination of differentially expressed genes were performed as described above and determined which CUGs and CDGs were altered by these mutations.

### Kaplan-Meier overall survival, Cox regression and ‘good’ versus ‘bad’ prognosis data analyses

Kaplan-Meier (KM) overall survival was performed for each gene in TCGA (146 tumors), GSE71729 (125 tumors), Puleo (288 tumor) and ICGC-AU (267 tumors) datasets. For TCGA, we selected the samples that were used for survival analysis in the original article on PDAC^4^. For the other datasets, we included all samples for which there is an accompanying survival data. We derived median expression value per gene and used it to rank samples into ‘high’ and ‘low’ groups for that gene. We then performed KM OS analysis with log-ranked test. Univariate Cox proportional hazards regression analysis was performed with the same sample size as used for KM except that the gene expression was set as continuous variables. The associated *P*-values was derived using the Wald statistic. Both the KM and Cox regression analysis were performed using *survival* package (v 2.43-3) in R and statistical significance was considered as *P*<0.05. Although all genes that were significantly predictive at least in one cohort were highlighted, we placed greater emphasis on genes that were significant in two or more cohorts. We also compared the consistent gene expression in tumors of patients with ‘bad’ prognosis (shorter survival time) vs ‘good’ (longer survival time) using the microarray dataset GSE42952 (tumors: *n*=6 good prognosis vs *n*=6 bad prognosis). The data was analyzed following the same procedure as in the five microarrays used for CUGs/CDGs identification.

### Receiver operating characteristics

We calculated the sensitivity (true positive rate) of each gene in TCGA and the Puleo data with respect to predicting tumor grade. For TCGA we used Grade 3 (*n*=69) and Grade 2 (*n*= 75) tumors, which constituted the entire data sample size. For Puleo data we analyzed Grade 3 (*n*=48) and Grade 2 (*n*=134), and in addition mainly the Grade 2 and Grade 1 (*n*=117). In each case, we performed receiver operating characteristics using the *pROC* package (v 1.13.0) and extracted the area under the curve (AUC) values. We used AUC > 60% as minimum cut off for sensitivity. Besides this, differential expression of genes in Grade 3 vs 2 in both TCGA and Puleo datasets were performed as described above for these datasets.

### Pathway and gene ontology analyses

Pathway analysis were performed using DAVID functional annotation platform (https://david.ncifcrf.gov/, v 6.8), or the gene set enrichment analysis tool (GSEA, v 4.0.3) with GSEAPreranked option. Ranking of the genes was based on the product of the logFC and -log(*P*-value) and analysis was based on only genes that were already determined to be significantly altered in the respective comparisons performed. GSEA for KEGG pathway, hallmark and transcriptional signatures were run with default parameters, except gene set size filter set at min=10. Gene ontology analyses were performed with DAVID. Details about the number of genes used are indicated in the Figure legends, where applicable.

### Assignment of genes to functional classes

We assigned the genes to their known functions, e.g., ‘metabolism’, ‘immunity’, etc, using published gene lists or gene lists extracted from KEGG or Molecular Signatures Database (MSigDB). For metabolism, we adapted a list of previously published 2,764 metabolic genes ^36^ and assorted them into metabolic and transporter genes. We also used a published list of 1,988 genes encoding transcriptional regulators in human^37^. We compiled a list of signaling genes (2,091 genes – KEGG), and a list of ~400 immune gene signatures from the Molecular Signatures Database (MSigDB)^38, 39^. To the extent possible, these genes were used, especially in Fig. 2, to assign the consistent genes to their known pathways.

### Determination of essential consistent genes

For essential gene analysis, we focused on the CUGs. To determine CUGs that are essential for PDAC viability, we used the CRISPR/Cas9 knockout screen data from Project Achilles, GECKO, Behan et al data assessed via BioGRID Open Repository of CRISPR Screens (ORCS)^1.0^ each of which covered >17,000 genes. In addition, we used shRNA knockdown screen data from Project Drive (includes ~8,000 genes). For Achilles we analyzed 24 PDAC cell lines with complete dependency scores; for GECKO we analyzed 8 cell lines, for ORCS it was 20 cell lines, and for Project Drive we analyzed 24 cell lines. Coverage of the CUGs was near complete. Specifically, of the 2,010 CUGs, we found 1,962 (~98%) in the Achilles data, 1,947 (~97%) in the GECKO data and 1,916 (95.3%) in Behan et al. ORCS data. The Project Drive, which was originally based on 7,975 genes predicted to be essential, included 972 CUGs (48%). We used the available CUGs per dataset to perform unsupervised hierarchical clustering. We then partitioned each dataset into two clusters: genes that their knockdown strongly impacted viability and genes with no strong impact. Only the CUGs that strongly impacted viability in at least three datasets were presented as essential/priority targets for PDAC.

### Cell line stratification and prediction of drug response

The 185 high priority therapeutic targets identified in this work was used to stratify PDAC cell lines in the cancer cell line encyclopedia (CCLE) (*n*=44 cell lines) and GSE57083 (AstraZeneca) datasets (*n*=23 cell lines). In the case of GSE57083, most cell lines existed as biological duplicates and for these their gene expression profiles were averaged to generate one datapoint per cell line as in the CCLE data. Thereafter, cell lines in the two datasets were stratified into those that have a high expression of the priority genes and those with low expression using clustering analysis. Following the identification of representative cells in high-expressing or low-expressing subgroups, the therapeutic sensitivity of the high-expressing groups were then determined using in the Genomics of Drug Sensitivity in Cancer (GDSC) data^23^, which contains experimentally determined sensitivity score for >195 compounds. Compounds that were effective against cell lines expressing a higher level of the high priority genes were selected for further validation studies.

### Cell culture, inhibitors, metabolomics

The human PDAC cell lines namely, BxPC-3, PA-TU-8902, ASPC1, CFPAC-1 and PANC1 were obtained from the American Type Culture Collection or the German Collection of Microorganisms and Cell Cultures (DSMZ). The mouse PDAC cell line used is the 7940b cells (C57BL/6J strain)^40^ derived from KPC tumor (*Ptf1a-Cre;LSL-Kra^sG12D^; Trp53^flox/+^*)–also called the KPC cell line. The cell lines were mycoplasma-tested (Lonza MycoAlert Plus, LT07-710), used within 10 passages and cultured in DMEM (Gibco, 11965-092) with 10% fetal bovine serum (FBS). The cells were cultured in 37°C incubator under humidified atmosphere. For the inhibitors tested: AZD5153 and gemcitabine were obtained from Cayman Chemicals (USA) while bromodomain containing protein inhibitor BD-9136 was a kind gift from Prof. Shaomeng. Wang (University of Michigan). Cell viability experiments were performed using CellTiter-Glo® 2.0 Cell Viability Assay (Cat # G9241, Promega) according to manufacturer’s instruction. Metabolomics profiling was performed by liquid chromatography tandem mass spectrometry (LC/MS/MS) and analyzed as previously described^26^.

### Subcutaneous xenograft tumor implantation

All animal studies were performed in accordance with the guidelines of Institutional Animal Care and Use Committee (IACUC) and approved protocol. For mouse tumor xenograft, ~6 weeks old female athymic nude mice NU/J (Stock No: 002019, The Jackson Laboratory) were maintained in the facilities of the Unit for Laboratory Animal Medicine (ULAM) under specific pathogen-free conditions. On the day of xenograft tumor injection, PA-TU-8902 cell line was harvested from culture plate according to normal cell culture procedures. The cells were counted, washed 1x with PBS and resuspended in 1:1 solution of serum free DMEM (Gibco, 11965-092) and Matrigel (Corning, 354234). Mice were subcutaneously (s.c.) injected on both flanks with 500,000 cell lines in 100 μL volume. When tumors became palpable, mice were randomized into the four treatment groups: vehicle, gemcitabine, BD-9136 and gemcitabine+BD-9136 following the same dosing regimen used for orthotopic experiment. Tumor size was assessed twice/week using a digital caliper and tumor volume was calculated as V = 1/2(length × width^2^). At endpoint, mice were euthanized, and the tumors harvested, weighed, and processed for further analysis.

### Mouse pancreatic orthotopic tumor model

To establish the orthotopic model, 50,000 KPC cells were injected into the pancreas of the wild type C57BL/6J mice (~8 weeks old). The cell lines were suspended at 1:1 ratio of serum free DMEM and Matrigel and injected in a 50 μL final volume. Five days post-injection, the mice were randomized into experimental and control groups (*n*=6 mice) based on body weight. Experimental groups consisted of gemcitabine (100 mg/kg body weight twice a week), BRD4 inhibitor BD-9136 (20 mg/kg body weight 5 days a week), or both. Vehicle consisted of 20% PEG400, 6% Cremophor EL, and ~74% PBS solution. Treatments were administered by intraperitoneal injection. At endpoint, body and tumor weight were taken and tissues extracted from further processing.

### Histology and Ki67 staining

Tumor tissue section and Ki-67 staining was done according to protocol in our lab as previously published^35,41^.

### Statistics and additional analysis

Details about the analyzed or plotted samples are described above and presented in Figure legends where applicable. Survival plots were generated with *survminer* package (v 0.4.3). Additional R packages used: *pheatmap* (v 1.0.12) and gplots (v 3.0.1) for heatmap, *ggfortify* for principal component analysis.

Network visualization and disease ontology analysis were performed with enrichplot (v 1.6.1) and DOSE (v 3.12.0) in R (v 3.6.2). We determined which CUGs or CDGs are well-studied or novel in PDAC by querying PubMed using each gene symbol. Search terms were the gene symbol and ‘pancreatic’ or ‘pancreatic cancer’. Genes that returned 0 or to less than 5 relevant results were considered novel.

## Data source and availability

All data used in this study are publicly available under the indicated accession numbers along with the clinical data where applicable. The microarrays were obtained from NCBI GEO or ArrayExpress databases. The ICGC-AU microarray data (release_28) was downloaded from https://dcc.icgc.org/projects/ along with the associated clinical data and had no embargo (March 2020). The cancer essential gene data from the Project Achilles, Project Drive and GECKO screens were downloaded from the DepMap Public 20Q1 (https://depmap.org/portal/download/), while ‘03_scaledbayesianfactor’ data from Behan et al^20^ CRISPR screen was downloaded from BioGRID ORCS (https://orcs.thebiogrid.org/Dataset/114). Drug response data were accessed via the Genomics of Drug Sensitivity in Cancer (GDSC) portal (https://www.cancerrxgene.org/).

## URLs

ArrayExpress database http://www.ebi.ac.uk/arrayexpress

cBioPortal https://www.cbioportal.org/

Human Protein Atlas https://www.proteinatlas.org/

International Cancer Genome Consortium (ICGC) data portal https://dcc.icgc.org/projects/

NCBI GEO Datasets: https://www.ncbi.nlm.nih.gov/geo/

DepMap https://depmap.org/portal/download/

## Supplementary Note

### 1. Additional independent cohorts

The additional datasets were included to ascertain whether any randomly selected PDAC cohorts will reflect the consistent genes and their expression pattern as found in the original five microarray datasets used for discovering the consistent genes. As shown in **Fig. 1c**, many CUGs and CDGs retained their expression pattern in these additional cohorts. Importantly, several topmost consistent genes also appeared within the top differential genes in each of the additional cohorts (**Extended Data Fig. 1c**). Thus, the identified consistent genes are reproducibly changed in most PDAC cohorts.

### 2. Consistency of gene expression in peripheral blood mononuclear cells

We analyzed three published microarrays of peripheral blood mononuclear cells (PBMCs) (**Extended Data Fig. 1a)**. We identified 672 genes consistently upregulated at least in two cohorts of PDAC PBMCs, including 54 genes in all three datasets (**Supplementary Table 4b**). In contrast, there were 920 consistently downregulated genes at least in two PBMC datasets, including 77 downregulated in all three (**Supplementary Table 4b**). These expressions reflect a less consistent gene expression pattern in blood compared to tissues, likely due to expected hemodynamic changes that are not applicable in tumors. Of the consistent genes, those downregulated in tissues (i.e., the CDGs) were particularly less overlapping. Specifically, only ~10% of the overlapping consistently downregulated genes in PBMCs showed same expression pattern with CDGs. Pathway alterations consistently underlying differentially expressed genes in the blood of PDAC patients are unknown. To this end we performed pathway annotation and GO analysis with all genes consistently expressed in at least two of the three PBMCs datasets analyzed. The result revealed alterations in immunologic and coagulation pathways and are presented in **Supplementary Table 4c**.

### 3. Expression pattern of the consistent genes in basal-like vs classical subtypes, proliferation-high, metastasized tumors and survival profile

Stratification of tumor samples (in TCGA, Moffitt, and Puleo cohorts) by basal-like and classical subtypes yielded a highly reproducible set of genes, including 889 genes (e.g., *A2ML1, KRT6A, MUC16, LY6D, KRT5, S100A2, KRT14*) highly expressed in basal-like tumors and 741 highly expressed genes (e.g., *REG4, CLCA1, TM4SF20, BTNL8, ITLN1, MYO1A, CLDN18*) in classical subtypes in all three patients’ cohorts. Similar reproducibility was observed with proliferation markers, where 956 genes were highly expressed and 731 lowly expressed in all three cohorts. Moreover, of the used proliferation markers, 8 (30%), i.e., *TOP2A, FOXM1, CDK1, CCNB1, RRM2, MKI67, BUB1*, and *TYMS* emerged in the top ranked 25 upregulated genes across the three cohorts (shown **Fig. 4b**). These data confirm the reproducibility of the stratification in these independent cohorts and reveal genes underlying proliferation in PDAC. There were genes that however showed discordant expression pattern. For example, *PSAT1, C9orf40, GPR3, PRPS2, SAMD1, IPO4, GIT1* and *DFFA* showed high expression in proliferation-high tumors but are consistently low in PDAC (i.e., CDGs). Such deviant expression pattern was also seen for 37 CUGs, notably including novel genes such as *IGFBP7, ISLR, ITGBL1, SFRP2, COL16A1, MATN3, PLXDC1*, and known genes such as *LYZ, TIMP1, NNMT, CXCR4*. We conclude that for these genes their expression pattern in tissues is not a reliable indices of tumor proliferation state.

With respect to metastasis, some CUGs were switched to ‘downregulated’ in metastasized tumors, while some CDGs are highly expressed. The switched expression pattern of these genes either suggest a less critical role in metastasis or may underscore our earlier point that at least more than two subtypes of PDAC exist given that several genes could not be easily assigned in a dichotomous analysis. There were also some inconsistencies in the pattern of survival prediction. For example, the high expression of glycolysis gene *ALDOA* predicted poor OS in Moffitt cohort but not in TCGA, whereas high *LDHA* predicted in TCGA but not Moffitt cohort. Further, high expression of zinc transporter *SLC39A10* predicted poor OS in TCGA, but better OS in the Moffitt cohort. Such molecular distinction could be leveraged upon for defining unique tumor subsets for functional characterization of highly context-dependent genes.

To determine expression changes at the initial progression phase, we analyzed a dataset on progressive PDAC stages, from intraductal papillary mucinous adenoma, neoplasia to carcinoma tissues^18^. Whereas 281 (14%) of the CUGs appeared at all three stages, most of the CUGs were expressed at the IPMA stages, further supporting a possible role of these genes at the tumor initiation and progression phases. Another strategy that could reveal genes that made may be involved in tumor progression is to examine genes that are differentially expressed across tumor grades. Comparison of Grade 3 versus 2 tumors in TCGA and Puleo cohorts revealed 309 differential genes (*P*<0.05), of which 178 were high and 131 were low. Genes in the respective tumor grade comparison are in **Supplementary Table 8**.

## Extended Data Figure Legends

**Figure S1. Consistently expressed genes in human PDAC tissues**

**a**, Gene datasets analyzed in the quest for consistent genes, priority targets and drug response predictors. In red are cell line datasets; GSE15932, GSE49641 and GSE74629 are microarray datasets of peripheral blood mononuclear cells. All other datasets are expression profile data from tissues.

**b**, The five ‘discovery’ microarray datasets of pancreatic ductal adenocarcinoma (PDAC) used for the identification of the consistently upregulated genes (CUGs) and consistently downregulated genes (CDGs). Genes were defined as ‘consistent’ if significantly upregulated or downregulated in at least 4 of the 5 datasets.

**c**, Heatmap showing that the additional microarray datasets reproducibly depicted the topmost consistent genes derived from the 5 ‘discovery’ datasets.

**d**, Topmost up- or downregulated genes that were expressed in 3 of the 5 datasets. In Figure S1c-d: orange – upregulated; blue – downregulated; blank/white – no data or not significantly changed in the corresponding dataset.

**e**, GSEA plots showing additional pathway enrichment in PDAC as derived with the consistent genes. Several genes such as *LAMC2, LAMB3* and genes encoding collagens are shared between ‘Focal adhesion’ and the ECM receptor interaction shown in Fig. 1e.

**f**, Interactome network analysis derived using the top 500 CUGs and 500 CDGs.

**g**, Enrichment plot showing the disease ontology associated with the top 500 CUGs and 500 CDGs. As shown, these genes are associated with various cancer types as well as premalignant lesions.

**Figure S2. Multiple molecular pathways underlie PDAC**

**a**, GSEA plot showing immune signatures upregulated of PDAC. The listed genes are top in the respective pathways.

**b**, Serine pathway genes are consistently downregulated in PDAC. *PSPH* was not significantly changed in any of the 5 datasets.

**c**, GSEA plot showing the downregulation of oxidative phosphorylation (OXPHOS); on the right, transsulfuration pathway and tryptophan pathways respectively.

**d**, GSEA plots showing the downregulation of AR and HNF1 signatures.

**e**, Heatmap showing epigenetic genes consistently upregulated or downregulated in PDAC.

**Figure S3. Consistent genes show tumor-specificity and expression changes in patients’ blood**

**a**, Heatmap showing CUGs that are the top lowly expressed or CDGs that are the top highly expressed in normal pancreas relative to other tissues from the Human Protein Atlas RNA seq data.

**b**, Heatmap showing CUGs that are the top lowly expressed or CDGs that are the top highly expressed in normal pancreas relative to other tissues from the genotype-tissue expression (GTEx) project.

**c**, Heatmap showing topmost CUGs lowly expressed or CDGs highly expressed in normal pancreas compared to normal liver, lymph, lung tissues in the GSE71729 (*P*<0.01).

**d**, Dot plot showing gene markers of various cell populations in the PDAC single cell RNA sequencing (scRNA seq) data used to validate tumor-specific and microenvironmental cell expression pattern of CUGs and CDGs.

**e**, UMAP plots visualizing T-cell (CD3E), myeloid (CD14) and acinar cell population in the scRNA seq data.

**Figure S4. The consistent genes correlate with tumor subtypes and aggressive features**

**a**, Schematic of analysis performed to identify association with clinicopathological variables. Tumor stratification was performed with TCGA (*n*=150 samples), GSE71729 (*n*=125), and Puleo (*n*=309) cohorts; mutation analysis was with TCGA, while OS and Cox regression was with TCGA (*n*=146), GSE71729 (*n*=125), Puleo (*n*=288) and ICGC-AU (*n*=269) cohorts.

**b**, Heatmap showing the stratification of three patient cohorts into classical and basal-like subtypes using the top 50 gene signatures published by Moffitt et al. (2015). Most of the 50 genes were captured in TCGA and Puleo et al. dataset.

**c**, Venn diagram showing the overlap of CUGs and CDGs with genes that were not significantly changed (*P*>0.05) in the comparison of basal-like and classical TCGA, Puleo et al. and GSE71729 (Moffitt) tumors. In bold indicates the number of CUGs/CDGs that are not changed in any of the three datasets. The topmost CUGs and CDGs are listed.

**d**, Representative patient single cell RNA seq data showing the epithelial cell (marked by KRT18/19) expression of basal-like, classical, and non-basal-like non-classical subtypes of PDAC.

**e**, Plots showing consistent genes in the topmost genes in metastasis vs normal primary tissues or metastasis vs primary tumors.

**f**, Venn diagram showing the overlap of CUGs and CDGs with genes differentially expressed in liver and peritoneal metastasis samples from the dataset GSE42952. Next, heatmaps showing expression of genes that overlapped between liver and peritoneal metastasis. The samples for liver or peritoneal metastasis were compared to samples of patients who had ‘bad’ prognosis.

**g**, Venn diagram showing the overlap of the basal-like/classical subtype signatures with liver metastasis signatures derived from GSE71729 and GSE19279 (liver metastasis vs liver normal tissues).

**h**, Venn diagram showing the overlap of GUGs and CDGs respectively expressed in various KRAS mutation tumors relative to tumors without KRAS mutation.

**i**, Pathways represented by genes upregulated or downregulated in tumors in KRAS G12D mutation.

**i**, Pathways upregulated or downregulated in Grade II relative to Grade I tumors from Puleo et al. datasets.

**k**, Heatmap showing top CUGs differentially expressed in premalignant pancreatic diseases, e.g., IPMA – intraductal papillary mucinous adenoma, IPMC – intraductal papillary mucinous carcinoma, IPMN – intraductal papillary mucinous neoplasms.

**l**, Venn diagram showing the number of CUGs and CDGs that are highly lowly expressed in tumors from patients that had ‘bad’ prognosis (pr.) (shorter survival time, <7 months, *n*=6) compared to those that had ‘good’ prognosis (longer survival time, >50 months, *n*=6) based on the dataset GSE42952.

**Figure S5. The high priority therapeutic targets predict epigenetic inhibitors for PDAC therapy.**

**a**, Schematic illustration of the cell line stratification. The 185 high priority targets were used to stratify PDAC cells in the cancer cell line encyclopedia (*n*=44 cell lines) and GSE57083 datasets (*n*=23 cell lines) into those expressing the priority genes at high and low levels, respectively.

**b**, Viability assay of PDAC cell lines treated with BETi AZD5153 alone or in combination with gemcitabine for 72 h.

**c-d,** Metabolomics profiling showing the metabolites altered in the intracellular compartment and culture media(extracellular) upon treatment with AZD5153 alone or in combination with gemcitabine for 24 h. **d**, Amino acid changes in media (extracellular compartment) of the treated TU8902 cell lines.

**e**, Tumor volume of the TU8902 xenograft treated with gemcitabine or in combination with BD-9136.

## Acknowledgments

We thank members of the Lyssiotis and Pasca di Magliano labs and the entire Pancreatic Disease Initiative at the Rogel Cancer Center, University of Michigan, for their insightful comments and discussions.

## Authors’ Contributions

Z.C.N conceived and designed the study, performed data analyses, experiments, and wrote the manuscript. H.G, M.N, V.C, D.K, S.K, N.S – data analysis; R.E.M, J.H, B.H – experiments; S.W, M.P, C.A.L – provided materials/reagents and designed experiments; M.P and C.A.L supervised the project. All authors read and approved the final manuscript.

## Abbreviations

CUGs: Consistently upregulated genes
CDGs: Consistently downregulated genes
GSEA: Gene Set Enrichment Analysis
GO: Gene ontology
KM: Kaplan-Meier
OS: Overall survival
OXPHOS: Oxidative phosphorylation
PDAC: Pancreatic adenocarcinoma
TCGA: The Cancer Genome Atlas
ICGC: International Cancer Genome Consortium

