## Supplementary Figures for "Multi-dimensional analyses identify genes of high priority for pancreatic cancer research"

### Extended Data 1.

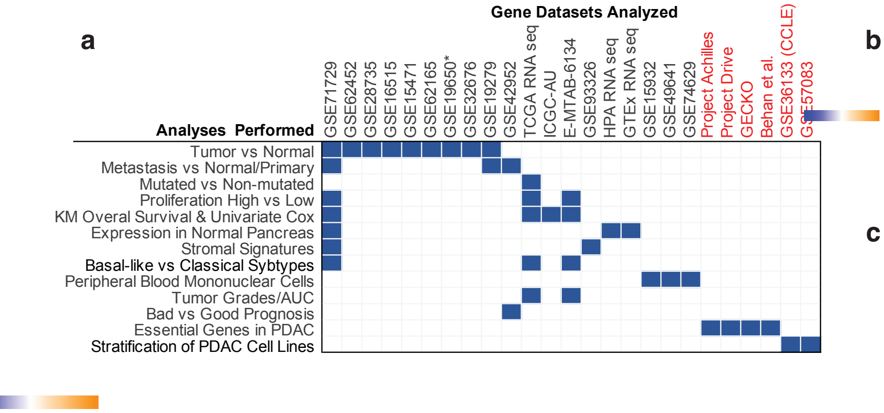


**b**

The 5 Discovery Datasets

| Microarrays | Non-Tumor | PDAC |
|---|---|---|
| GSE71729 | 46 | 145 |
| GSE62452 | 61 | 69 |
| GSE28735 | 45 | 45 |
| GSE16515 | 16 | 36 |
| GSE15471 | 36 | 36 |
| Total | **204** | **331** |

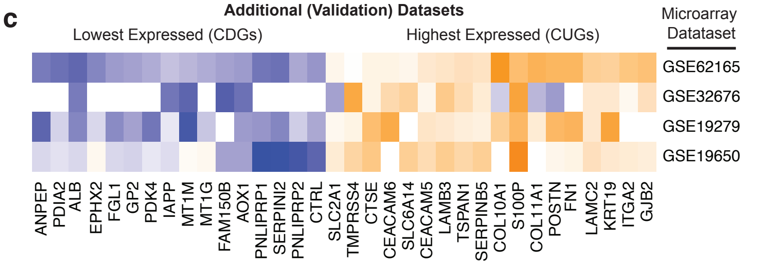


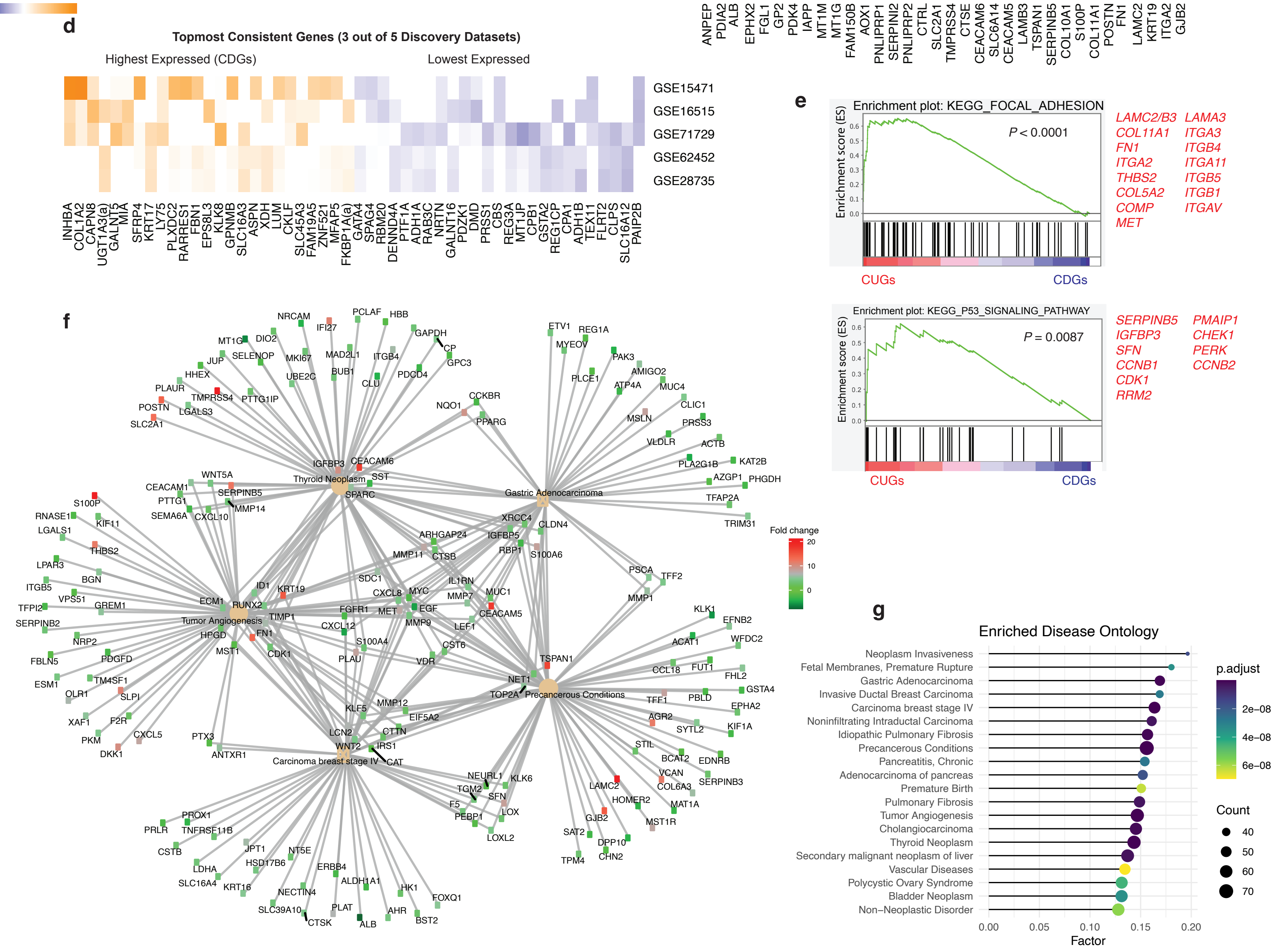

#### Extended Data 2.

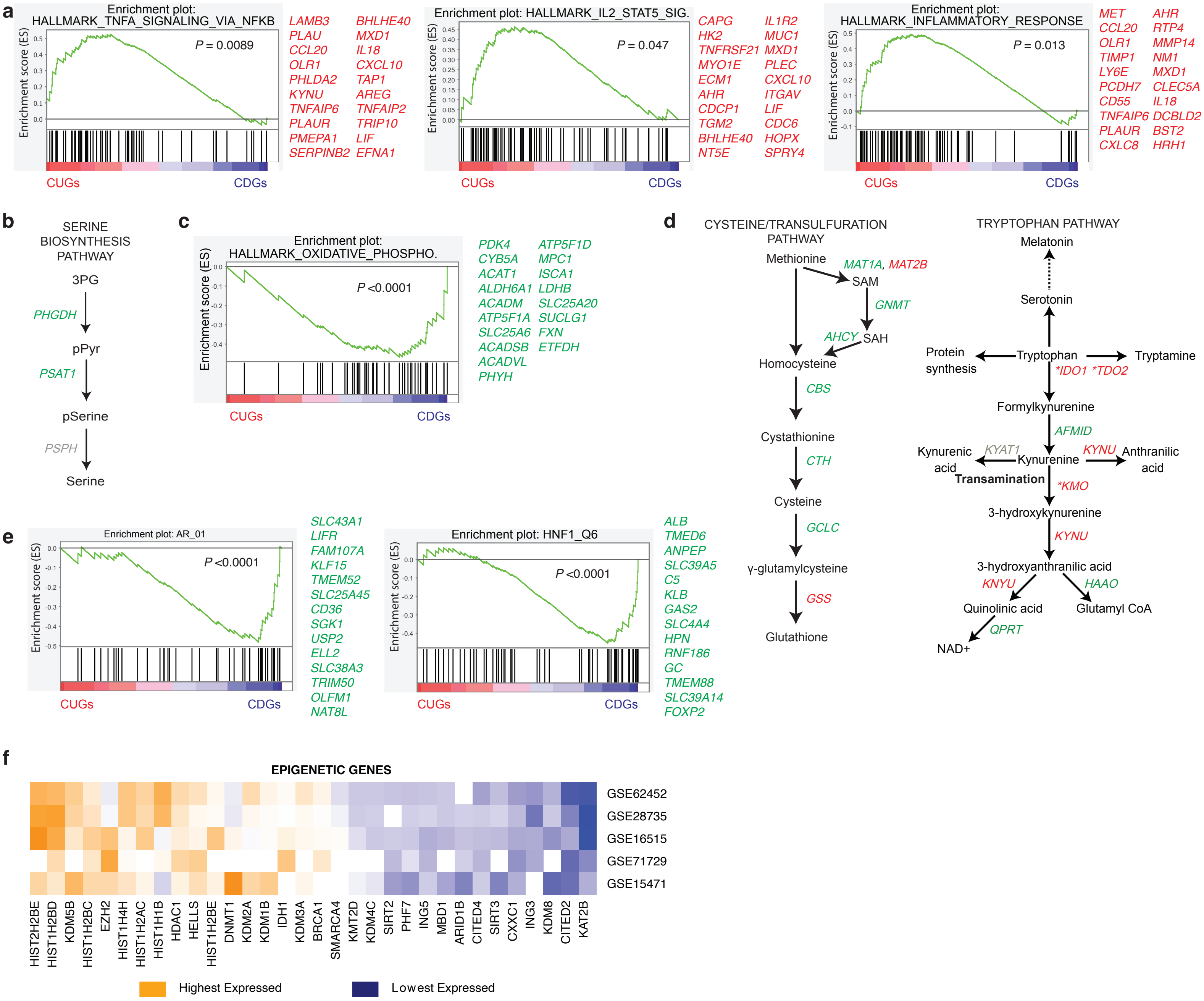

### Extended Data 3.

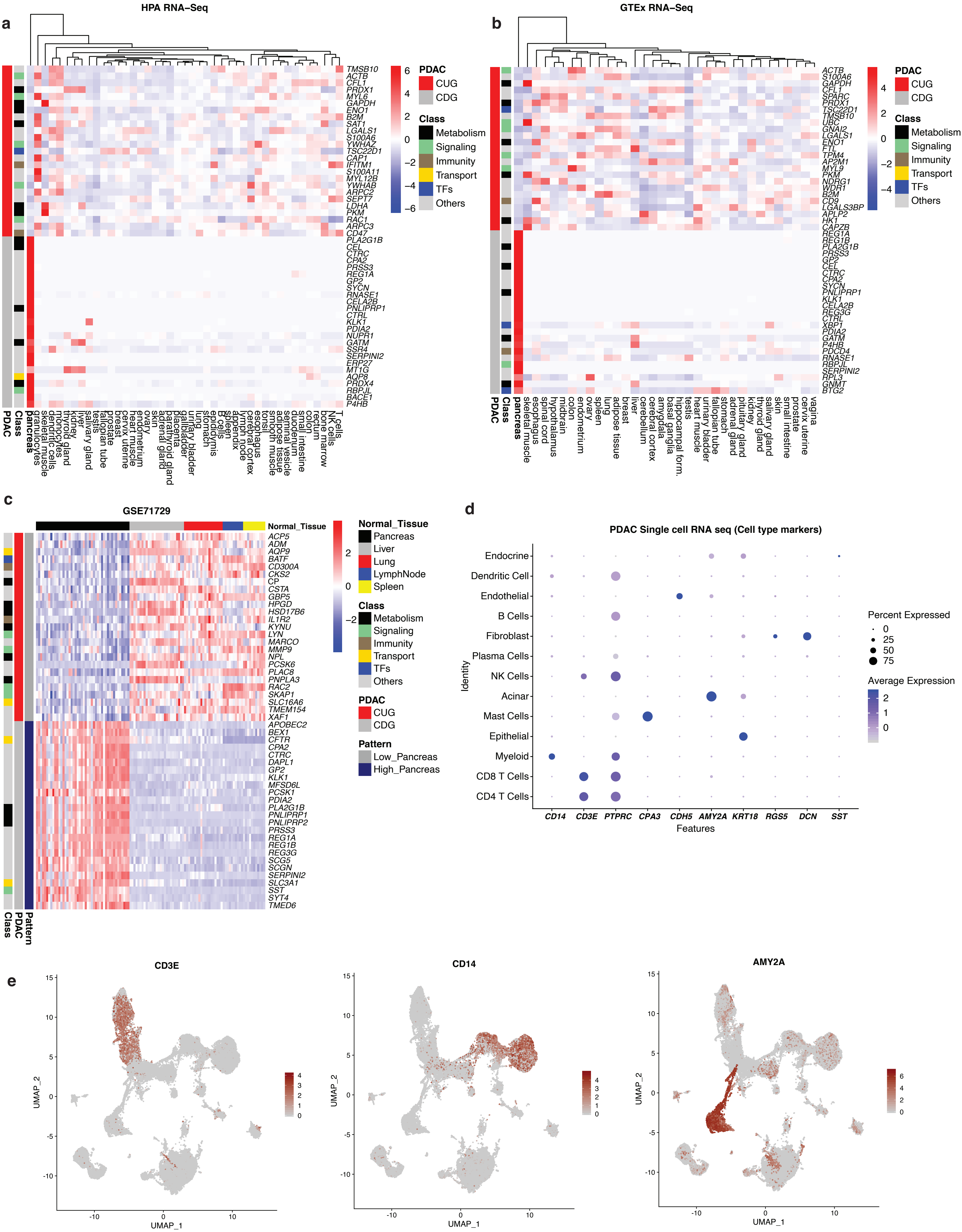

### Extended Data 4.

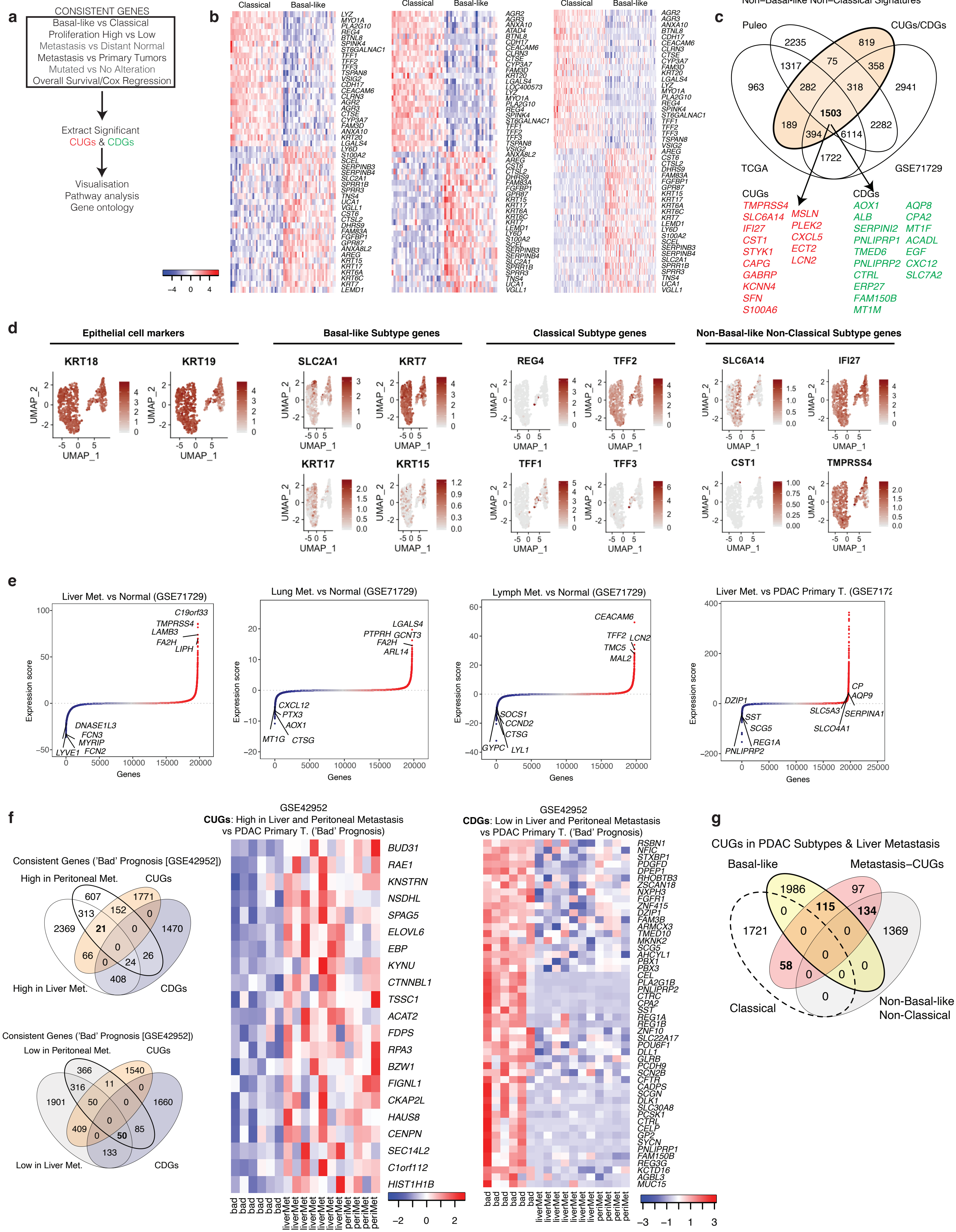

**Extended Data 4.**

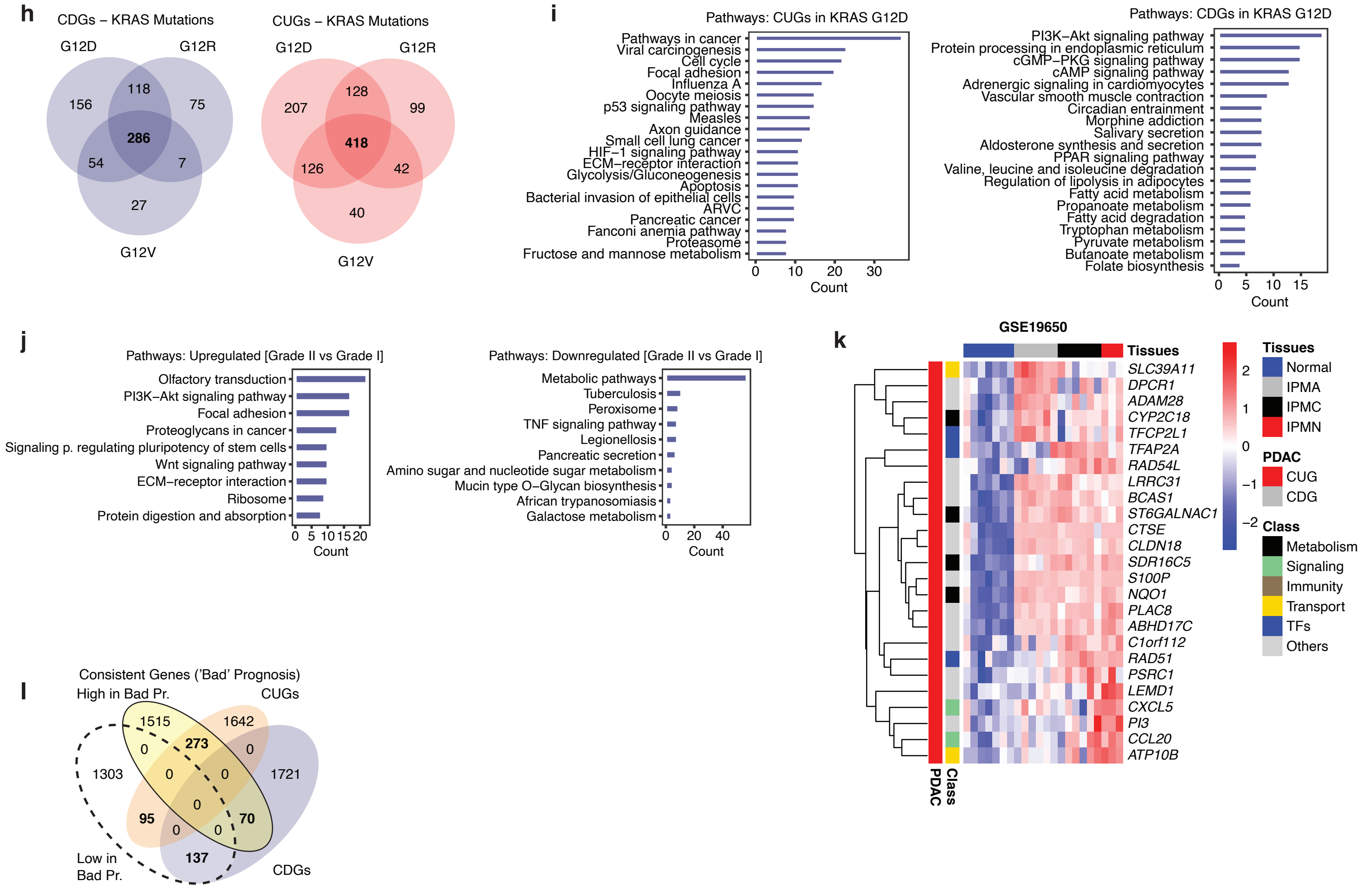

#### Extended Data 5,

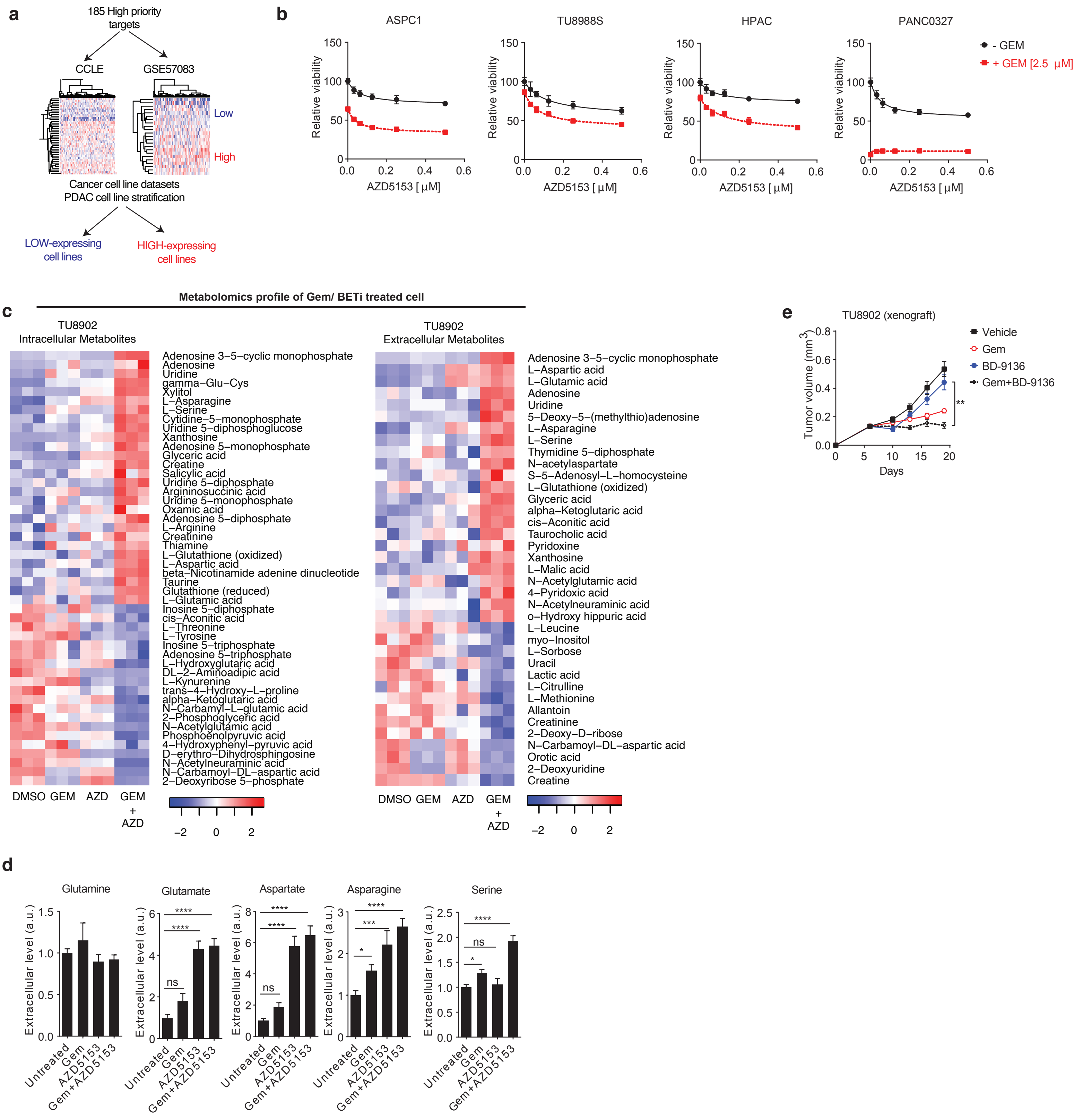
